# Characterisation of novel *Campylobacter jejuni* Type VI secretion system (T6SS) effectors and exploration of the roles of the *C. jejuni* T6SS in bacterial antagonism and human host cell interaction

**DOI:** 10.64898/2026.03.25.714310

**Authors:** Zahra Omole, Subhadeep Gupta, Molly Webster, Janie Liaw, Geunhye Hong, Cadi Davies, Abdi Elmi, Nicolae Corcionivoshi, Brendan W. Wren, Ezra Aksoy, Daniel Inaoka, Amirul Islam Mallick, Abderrahman Hachani, Nick Dorrell, Ozan Gundogdu

## Abstract

*Campylobacter jejuni* is a leading global cause of acute foodborne gastroenteritis however, *C. jejuni* lacks some of the classic virulence determinants associated with other common enteric bacterial pathogens. In recent years an increasing number of *C. jejuni* isolates have been identified to encode Type Six Secretion System (T6SS), an apparatus utilised by Gram-negative bacteria to secrete toxic bacterial effectors into neighbouring cells. Despite the prevalence of the T6SS and previous investigations, the roles of the *C. jejuni* T6SS are still not well characterised especially when compared to our knowledge of other clinically relevant T6SS-positive bacterial species. Additionally, as of yet, no *C. jejuni* T6SS cargo effectors have been characterised. In this study, we show the *C. jejuni* 488 strain T6SS displays contact-dependent antagonistic behaviour towards T6SS-negative *C. jejuni*, *Campylobacter coli*, *Escherichia coli* and *Enterococcus faecium* strains suggesting the presence of the T6SS contributes to the competitive capacity of this *C. jejuni* T6SS-positive strain. Moreover, this antagonistic activity is linked to the functionality of CJ488_0980 and CJ488_0982, two novel putative Tox-REase-7 domain-containing effectors, which were identified through bioinformatical analysis of the *C. jejuni* 488 strain genome. Additionally, our investigations propose the *C. jejuni* 488 T6SS contributes to interaction, invasion and intracellular survival in human intestinal epithelial cells (IEC). Collectively, these initial findings are the first examples of *in vitro* investigation of putative cargo effectors in *Campylobacter* spp. and provide valuable insights into the roles of *C. jejuni* T6SS effectors in bacterial competition and pathogenesis. This study highlights the importance of T6SS as an emerging virulence determinant in *Campylobacter* spp. warranting further investigation.

## Introduction

Bacteria utilise transmembrane secretion machineries to translocate proteins, small molecules, and DNA across membranes to interact with the environment, competitors, and host cells (Costa et al., 2015, Filloux, 2022). The Type Six Secretion System (T6SS) is contractile nanomachine encoded by over 25% of Gram-negative bacteria which facilitates the secretion of toxic effector proteins into the extracellular milieu or directly into target eukaryotic or prokaryotic cells (Monjaras Feria and Valvano, 2020, Chang et al., 2014, Singh and Kumari, 2023). The T6SS machinery resembles an inverted bacteriophage tail, it can be simplified to a hollow needle-like structure (TssD), encased by a contractile sheath (TssBC), tipped by a puncturing (VgrG) spike which is sharpened by a PAAR-like (proline–alanine–alanine–arginine) protein (He et al., 2023). This is built on a baseplate (TssEFGK) anchored to the cell-spanning membrane complex (TssJLM) (Cherrak et al., 2019). Upon activation of the T6SS machinery, the sheath contracts which triggers the propulsion of the spiked inner needle outside the cell, allowing the delivery of recruited effectors into a target cell or the extracellular space (Brunet et al., 2015).

The role(s) of T6SS are inherently dependent on the functionality of the individual effectors and the diversity of the effector repertoire that are secreted through the apparatus (Monjaras Feria and Valvano, 2020, Jurenas and Journet, 2021). Effectors are typically classified depending on their method delivery as specialised or cargo effectors (Alcoforado Diniz et al., 2015). Specialised or “evolved” T6SS effectors possess *N*-terminal secreted canonical T6SS structural elements (e.g. TssD, PAAR and VgrG) and toxic *C*-terminal fused specialised extended regions granting effector activities (Kanarek et al., 2023, Geller et al., 2024). Whilst cargo effectors are toxic domain-containing proteins which lack a canonical secretion signal so must non-covalently bind with secreted T6SS machinery components, often aided by chaperons and adapter proteins, to facilitate T6SS-dependent secretion (Kanarek et al., 2023, Jurenas and Journet, 2021). The majority of characterised T6SS effectors possess antibacterial activities and are used to eliminate competitors present in polymicrobial environments (Zhao et al., 2018, Russell et al., 2014). However, a collection of T6SS effectors display anti-eukaryotic properties and subvert eukaryotic cell processes or modulate immune signalling to enhance virulence and survival in the host (Hachani et al., 2016, Pukatzki et al., 2006, Chen et al., 2017). More recently effectors with trans-kingdom functions and the ability to target fungi and Gram-positive bacteria have been described (Wang et al., 2025, Colautti et al., 2025, Trunk et al., 2018, Le et al., 2021).

*Campylobacter jejuni* is the leading global cause of foodborne bacterial gastroenteritis (Elmi et al., 2020). *C. jejuni* infections typically present with symptoms of abdominal pain, vomiting and watery or bloody diarrhoea (Kaakoush et al., 2015). *C. jejuni* infections are also implicated in the development of Guillain-Barré syndrome (GBS) and Miller Fisher syndrome (MFS), rare post-infectious severe neuropathological disorders (Nyati and Nyati, 2013, Ang et al., 2001). However, in low-resource settings where *C. jejuni* is endemic, recurrent infections are associated with malnutrition, growth impairment and fatality, as such the global economic and health burdens associated with *C. jejuni* infections are high (Amour et al., 2016, Zhang et al., 2022). Despite the prevalence and the global economic and health burdens associated with *C. jejuni*, the virulence determinants of *C. jejuni* are still poorly understood (Omole et al., 2024).

Prevalence studies have revealed T6SS-positive *C.jejuni* are highly represented by isolates belonging to the ST-353, ST-460, ST-607, and ST-446 clonal types (Rokney et al., 2018, Katz et al., 2023, Harrison et al., 2014, van Vliet et al., 2021). Host association studies have identified isolates in these phylogenetic lineages as chicken specialist isolates or lineages that are consistently associated with incidences of human gastroenteritis (Kelley et al., 2020, Dunn et al., 2018, Cody et al., 2012, van Vliet et al., 2021). With such a high proportion of T6SS-positive clinical isolates being restricted to specific lineages, the T6SS and its putative effectors are proposed to increase strain fitness within the gastrointestinal tract of humans and chickens (Katz et al., 2023). Phenotypic characterisation studies have ascribed the *C. jejuni* T6SS with roles including involvement in host colonisation, resistance to bile acids and oxidative stress, contact-dependent lysis of erythrocytes and bacterial antagonism (Liaw et al., 2019, Bleumink-Pluym et al., 2013, Gupta et al., 2021, Lertpiriyapong et al., 2012). However, these functions have yet to be associated with any specific effectors. Whilst *in silico* investigations have predicted the presence of several T6SS effectors candidates so far TssD, a structural component, remains the only secreted T6SS protein that has been investigated through *in vitro* studies (Lertpiriyapong et al., 2012, Katz et al., 2023, Robinson et al., 2021). As *tssD* knockout mutation renders the *C. jejuni* T6SS non-functional it has been difficult to unpick the function of TssD from the overall role of the T6SS (Liaw et al., 2019, Bleumink-Pluym et al., 2013).

In this study, we investigated the role of the *C. jejuni* T6SS in antibacterial competition and interaction with human host cells using the T6SS-positive *C. jejuni* 488 strain (ST-353) a Brazilian clinical isolate. Three novel putative effector proteins (CJ488_0980 and CJ488_0982) found encoded in the downstream genetic variable region on the T6SS gene encoding *Campylobacter jejuni* Pathogenicity Island 1 (CJPI-1 PAI) in *C. jejuni* 488 strain were characterised using phenotypic characterisation assays with defined isogenic mutants (488 *cj0980* and 488 *cj0982)*. Our findings propose CJ488_0980 and CJ488_0982 are bona-fide T6SS effectors involved in bacterial antagonism through contact-dependent killing of prey T6SS-negative bacteria. Additionally, the *C. jejuni* 488 strain T6SS enhances interaction and invasion of human host intestinal epithelial cells (IECs). The findings of this study have implications on the role of the T6SS *C. jejuni* as an emerging virulence determinant and suggest the presence of the *C. jejuni* T6SS increases fitness against competitors present within the human and chicken gut.

## Materials and Methods

### Bacterial strains and growth conditions

All *C. jejuni* and *C. coli* wild-type strains and mutants used in this study are listed in the supplementary (Table S1 and Table S2). *C. jejuni* and *C. coli* were grown at 37°C under microaerobic conditions (85% of N_2_, 5% of O_2_ and 10% of CO_2_) in a variable atmosphere chamber (Don Whitley Scientific, UK). *C. jejuni* and *C. coli* were grown on Columbia Blood Agar (CBA) plates (Oxoid, UK) supplemented with 7% (v/v) horse blood in Alsever’s (TCS Microbiology, UK) and *Campylobacter* selective supplement Skirrow (Oxoid, UK) or in *Brucella* broth (BD Diagnostics, UK) whilst shaking at 75 rpm. When appropriate, CBA plates and *Brucella* broth were supplemented with kanamycin (50 μg/ml), chloramphenicol (10 μg/ml) or tetracycline (20 µg/ml). *Enterococcus faecium* was grown on CBA plates supplemented with 7% (v/v) horse blood in Alsever’s and *Enterococcus faecium* Selective Supplement (Sigma-Aldrich, USA) or in MRS broth (De Man, Rogosa, Sharpe broth; Thermo Fisher Scientific, USA) whilst shaking at 75 rpm under microaerobic conditions at 37°C in the variable atmosphere chamber. The *Escherichia coli* strains used in this study are listed in supplementary (Table S2). *E. coli* were grown at 37°C under aerobic conditions on lysogeny broth (LB; Oxoid) agar plates or in LB broth shaking at 200 rpm (Sanyo, UK). When necessary, *E. coli* cultures and LB plates were supplemented with ampicillin (100 µg/ml), kanamycin (50 μg/ml) or chloramphenicol (50 µg/ml).

### Construction of the *C. jejuni* mutants

The *C. jejuni* 488 *cj0980* mutant and *C. jejuni* 488 *cj0982* mutant were constructed using splicing by overlap extension PCR (SOE PCR) and allelic homologous exchange to introduce a non-functional version of the gene disrupted by an antibiotic resistance cassette into the open reading frame (ORF). (Horton et al., 2013). Briefly for the construction of the *C. jejuni* 488 *cj0980* mutant, primers were designed to amplify the 5’ and 3’ fragment of the *cj0980* gene from *C. jejuni* 488 wild-type genomic DNA excising ≈ 200 bp from the centre of the ORF (Supplementary Table S3). The internal primers were constructed to be partially complementary to each other and introduce a BglII restriction site with the overlapping. The 3’ and 5’ fragment arms were then combined and amplified using SOE PCR with the external primers. The purified PCR products were ligated into a pGEM^®^-T Easy Vector (Promega, USA) and transformed into XL1-Blue Competent Cells (Agilent Technologies, UK). The resulting transformed plasmid was digested using the Bg1II restriction site to allow insertion of the BamHI-digested Kanamycin resistance (Kan^r^) cassette amplified from the pJMK30 vector. Insertion of the cassette in the correct orientation was confirmed by Sanger sequencing and the resulting plasmid was transformed into *C. jejuni* 488 wild-type strain by electroporation.

A vector-free methodology was used for the construction of the *C. jejuni* 488 *cj0982* mutant. This allowed a linearised copy of the *cj0982* gene disrupted by the Erythromycin resistance (Ery^r^) cassette, amplified from the pDH20 vector, to be directly transformed into *C. jejuni* 488 wild-type strain by electroporation. The fragments were constructed as previously described with the following exception, the internal primers were complementary to the Ery^r^ cassette which permitted the direct splicing of the Ery^r^ cassette with the 3’ and 5’ arm fragments of the 488 *cj0982* insert using SOE PCR (Supplementary Table S3). Successful construction of the *C. jejuni* 488 *cj0980* mutant and the *C. jejuni* 488 *cj0982* mutant were confirmed through PCR, sanger sequencing and whole genome sequencing.

*C. jejuni* 488 wild-type expressing green fluorescent protein (GFP) was constructed as previously described, with a pCJC1 harbouring *gfp* used to transform the wild-type strain through electroporation (Jervis et al., 2015).

### Real-Time Quantitative Polymerase Chain Reaction (qRT-PCR) Analyses

*C. jejuni* RNA was isolated from *C. jejuni Brucella* broth cultures or infected Human intestinal epithelial cells using PureLink^TM^ RNA mini kit (Thermo Fisher Scientific). Contaminating DNA was removed from the RNA samples using TURBO DNA-free kit (Ambion, USA). The total RNA concentration and purity was measured using a NanoDrop ND-1000 spectrophotometer (Thermo Fisher Scientific). 400 ng of each RNA sample was used for complementary (cDNA) synthesis using SuperScript III First-Strand Synthesis System (Thermo Fischer Scientific). qRT-PCR primers used in this study can be found in Supplementary Table S4. qRT-PCR was performed on an ABI-PRISM 7500 instrument (Applied Biosystems Life, USA), with reactions were performed with SYBR Green PCR Master Mix (Applied Biosystems) as previously described (Hong et al., 2025). Relative expression levels were compared using the comparative threshold cycle (2^-ΔΔCT^) method as previously outline with *rpoA* analysed and used to normalise expression levels of the target genes (Schmittgen and Livak, 2008).

### Whole cell lysate and supernatant preparation, SDS-PAGE and Western blot Analysis

*C. jejuni* harvested from 24-hour CBA plates were used to inoculate *Brucella* broth flasks to an OD_600_ (optical density at 600 nm) of 0.1, the cultures were incubated under microaerobic conditions whilst shaking at 75 rpm. The broth cultures were centrifuged at 4°C and 4,000 rpm for 30 minutes to separate the whole cell pellet and supernatant. The supernatant fraction was prepared by filtering through a 0.2 μm-pore-size filter (Merk Millipore, USA) and further concentrated using successive 30-minute centrifuge spins at 4,000 rpm at 4°C in a 10 kDa Amicon Ultra-15 centrifugal filter (Merck Millipore). The whole cell lysate was sonicated for 5 minutes on ice using 50% amplitude with a Model 120 Sonic Dismembrator (Thermo Fisher Scientific). The protein concentrations of the samples were measured using the Pierce^TM^ Bicinchoninic acid (BCA) Protein Assay kit (Thermo Fisher Scientific). Samples were diluted using DEPC-treated water (Invitrogen) and 2X Laemmli buffer (Sigma-Aldrich) to the desired concentration, incubated at 95°C for 10 minutes and centrifuged at 13,000 rpm for 5 minutes. 30 μg of the prepared whole cell lysate and 30 μg of the supernatant protein samples were separated using 4-12% (w/v) NuPAGE™ Bis-Tris gel (Thermo Fisher Scientific) in 1X MOPS running buffer (Invitrogen). After the run, the separated proteins were transferred onto a nitrocellulose membrane using iBlot® 2 transfer stacks (Life Technologies, USA) and iBlot® Gel Transfer Device (Life Technologies). The membranes were blocked with 1X phosphate-buffered saline (PBS; Thermo Fisher Scientific) with 2% (w/v) skimmed milk overnight at 4°C and then probed with primary antibodies as previously described (Liaw et al., 2019). The following primary antibodies were utilised in this study; TssD antibody (Capra Science Antibodies AB, Sweden) and CysM (*O*-acetylserine sulfhydrylase B) (Liaw et al., 2019, Christensen et al., 2009). After incubation the blots were developed using IRDye® 680RD goat anti-rabbit IgG Secondary Antibody (LI-COR Biosciences, USA) and imaged using the LI-COR Odyssey Classic (LI-COR Biosciences).

### Interbacterial competition assays

The *C. jejuni* 488 *gfp* strain was used as an attacker instead of the 488 wild-type strain to permit selection by the chloramphenicol resistance marker of the 488 *gfp* during competition assays. Interbacterial co-culture assays were performed as described by by Hachani *et al* with minor modifications (Hachani et al., 2013). Briefly 24 hour CBA plate cultures of *C. jejuni* and *C. coli* were individually harvested and adjusted to an OD_600_ of 0.5 in sterile PBS. Attacker and prey bacteria were mixed in a ratio of 1:1 or 5:1 with 25 μL of the resulting mixed bacterial suspensions plated in triplicate onto 1.5% (w/v) *Brucella* agar. Alongside this, unmixed control individual inputs of each attacker or prey were mixed in the same ratios with PBS and then spotted onto 1.5% *Brucella* agar. After 16 hours of incubation at 37°C under microaerobic conditions, bacteria was recovered in 1 ml PBS and serially diluted. The dilutions were plated simultaneously onto CBA supplemented with appropriate antibiotics to select for either the attacker or prey bacteria (See Table S1 and Table S2). After incubation for 48 hours at 37°C under microaerobic conditions, the recovered input and output colony-forming units (CFUs) were counted and used to calculate the competitive index values using the following formula:

(output attacker/output prey)/ (input attacker/input prey) = competitive index.

To measure reliance on contact for T6SS, once the individual suspensions of *C. jejuni* attacker and prey were prepared as previously described, 25 μl of the attacker *C. jejuni* was spotted directly on the 1.5% (w/v) *Brucella* agar, followed by a sterile 0.2 μm pore size membrane (Whatman, USA) and lastly 5 μl of the prey *C. jejuni* was spotted on top of the membrane. Plates were incubated and processed as previously described with loops used to harvest the membrane and bacteria which were all resuspended in 2.5 ml of PBS before serial dilutions were performed.

For interbacterial assays involving *E. faecium* or *E. coli* as prey the following modifications were made. An overnight culture *of E. faecium* grown under microaerobic conditions at 37°C in MRS broth was adjusted to an OD_600_ of 0.5 before being mixed with attacker *C. jejuni* in 5:1 ratio of attacker to prey in PBS. 25 μl of the resulting mixture was spotted on 1.5% (w/v) *Brucella* agar supplemented with 7% (v/v) horse blood in Alsever’s. The plates were then incubated and processed as previously described. For selection of *E. faecium,* serial dilutions were made on CBA plates supplemented with *Enterococcus faecium* Selective Supplement (Sigma-Aldrich) and the *E. faecium* colonies were counted after 24 hours with CFU/ml calculated and used to calculate the competitive index as before. For interbacterial competition assays involving prey *E. coli* DH5α, overnight *E. coli* LB cultures were adjusted to an OD_600_ of 0.5 in Mueller Hinton Broth (Oxoid). Attacker *C. jejuni* and prey *E. coli* were then mixed in a 5:1 or 50:1 ratio respectively, and 25 μl of the resultant suspension was spotted onto Mueller Hinton aga. All plates were then incubated under microaerobic conditions at 37°C for 16 hours. Spots were harvested with an inoculating loop, resuspended in PBS, and serially diluted. Concurrently each dilution was plated on *Campylobacter* Blood-Free Selective Agar Base (Thermo Fisher Scientific) plates supplemented with CCDA (Charcoal Cefoperozone Deoxycholate agar) Selective Supplement (Thermo Fisher Scientific) and the appropriate antibiotics to select for *C. jejuni* and Mueller Hinton agar supplemented with X-gal (40 μg/ml) and IPTG (0.1 mM) to select for the prey *E. coli* DH5α. Mueller Hinton agar plates were incubated overnight under microaerobic conditions at 37°C whilst CCDA plates were incubated under the same conditions for 48 hours. CFUs of the input and output attacker and prey bacteria were counted and used to calculate the competitive index as before.

### Human intestinal epithelial cell (IEC) lines and cell culture conditions

Human carcinoma cell lines T84 cells (ECACC 88021101) and Caco-2 cells (ECACC 86010202) obtained from the European Collection of Authenticated Cell Cultures (ECACC; UK) were used for this study. T84 and Caco-2 cells were cultured at 37°C in a 5% CO_2_ humidified atmosphere in complete growth medium, a 1:1 mixture of Dulbecco’s Modified Eagle’s Medium supplemented and Ham’s F-12 medium (DMEM/F-12; Thermo Fischer Scientific), supplemented with 1% (v/v) of MEM non-essential amino acids (Sigma-Aldrich), 1% (v/v) penicillin-streptomycin (Sigma-Aldrich) and 10% (v/v) of foetal bovine serum (FBS; Labtech, UK).

### interaction, invasion, and intracellular Survival assays

Interaction, invasion, and intracellular survival assays were performed using T84 and Caco-2 cells seeded in a 24-well plate as previously described (Hong et al., 2023). Before infection, T84 and Caco-2 cells were washed with PBS and the culture medium was replaced with complete growth medium without penicillin-streptomycin. The IECs were infected with *C. jejuni* with an OD_600_ of 0.2 (≈ 2 x 10^8^ CFU/ml) giving a multiplicity of infection (MOI) of 200:1 and incubated at 37°C in a 5% CO_2_ incubator. For interaction (adhesion and invasion) assays after 3 hours of infection with *C. jejuni*, the monolayers were washed three times with PBS and then lysed with 0.1% (v/v) Triton X-100 (Sigma-Aldrich) for 20 minutes at room temperature. The cell lysates were serially diluted with PBS, plated on CBA plates and incubated under microaerobic conditions at 37°C for 48 hours before CFU/ml representing the number of *C. jejuni* interacting with the cells were enumerated. For invasion assays, after 3 hours of *C. jejuni* infection, the IECs were treated with gentamicin (150 μg/ml) for 2 hours to kill the extracellular bacteria. The IECS were then washed, lysed and the cell lysates were serially diluted and plated on CBA plates. After 48 hours colonies were counted and the CFU/ml representing the invaded bacteria, was calculated. Lastly for the intracellular survival assay after 3 hours of infection with *C. jejuni*, the IECs were treated with gentamicin (150 μg/ml) for 2 hours to kill the extracellular bacteria, then an additional 18 hour incubation with gentamicin (10 μg/ml) was conducted. After incubation the IECs were washed, lysed, serial diluted and plated onto CBA plates as before.

### Statistical Analysis

All experiments were performed with at least three technical replicates conducted within each biological replicate and with a minimum of three biological replicates performed. Statistical analysis and graphing were performed using GraphPad Prism 8.1.2 Software (GraphPad software, USA). The mean between two independent data sets were compared for significance using unpaired *t*-test with * indicating *p* = < 0.05, ** indicating *p* = < 0.01, *** indicating *p* = < 0.001 and **** indicating *p* = < 0.0001.

## Results

### The *C. jejuni* 488 T6SS displays antagonism towards T6SS-negative *C. jejuni*

To investigate if the *C. jejuni* T6SS outcompetes non-kin bacterial strains and shapes inter-strain microbial dynamics, T6SS-negative strains; 81-176 and 11168H *kpsM* were used as prey during competitive index assays. Competition experiments relied on distinguishing between attacker and prey bacteria which was achieved using the inherent natural resistance or engineered antibiotic resistance of the bacteria to allow selection of the recovered output after co-incubation of bacteria. As such, 488 *gfp* was used for these experiments instead of 488 wild-type strain to permit attacker output enumeration through antibiotic selection. Growth kinetics and TssD secretion immunoblots revealed no significant differences between the 488 *gfp* strain and the 488 wild-type strain (Supplementary Fig. S1). The previously constructed 488 *tssD* mutant and 488 *tssD*+ complement were used as attackers to investigate the role of a non-functional and fully functional T6SS on competitive advantages. TssD is essential for assembly and activity as it forms the tubular structure that facilitates the delivery of the effectors into target cells, as such inactivation of *tssD* results in a non-functional T6SS (Liaw et al., 2019).

Inter-strain competitive index assays revealed *C. jejuni* 488 *gfp* displays antagonistic behaviour towards both prey T6SS-negative *C. jejuni* (81-176 strain and 11168H *KpsM* mutant). However, this phenotype was not exhibited by the 488 *tssD* mutant, which has a non-functional T6SS, during competition with prey 81-176 or during competition between the two T6SS-negative strains (Fig. 1). Notably the competitive advantage phenotype is exhibited during competition assays with the functional T6SS 488 *tssD+* complement*+* as attacker. This suggests that the functionality of the *C. jejuni* 488 T6SS is responsible for this predatory behaviour. The 488 *tssD* mutant, 488 *tssD+* complement and 11168H *kpsM* mutant possess the same antibiotic resistance marker which prevented competition assays from being conducted between these strains. Nevertheless, the same behaviour of antagonism was observed by the *C. jejuni* 488 *gfp* towards 11168H *kpsM* mutant. Competition assays also revealed that *C. jejuni* 488 T6SS-mediated antibacterial activity was density-dependent as the presence of attackers 488 *gfp* or 488 *tssD*+ complement at a higher cell density to prey, elicited greater prey killing (Supplementary Fig. S2).

**Figure 1:**
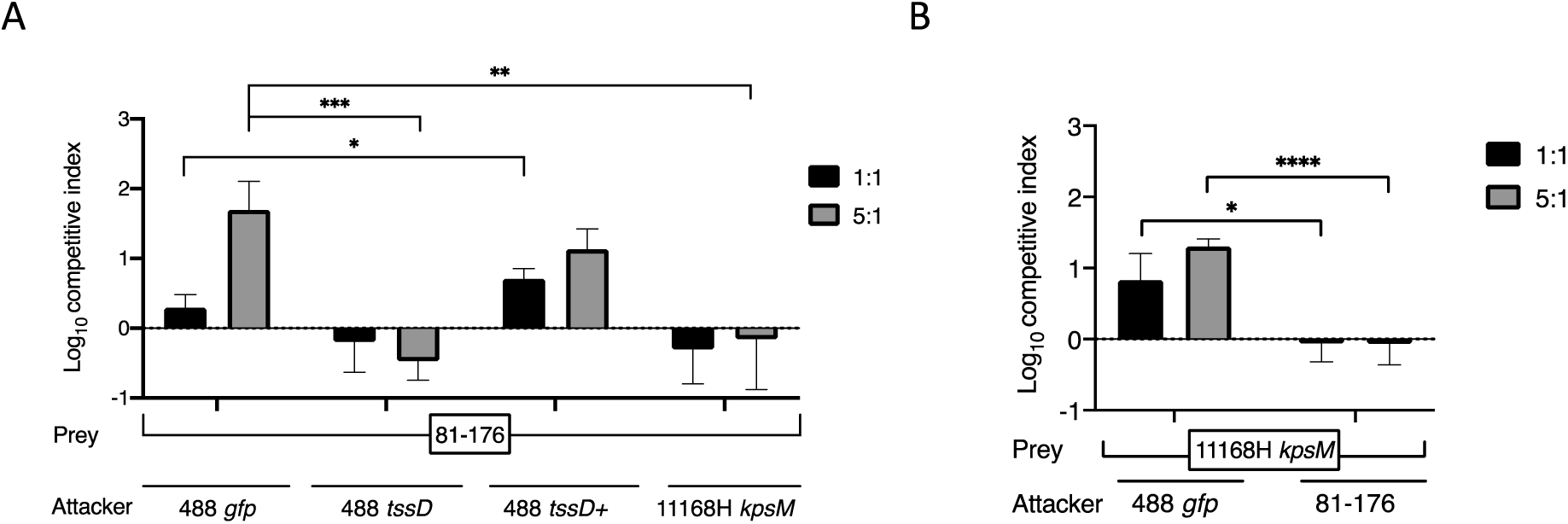
*C.jejuni* 488 T6SS targets T6SS-negative *C. jejuni* in a density-dependent manner. Competition assays were conducted between the indicated *C. jejuni* attacker and (A) prey 81-176 strain or (B) prey 11168H *kpsM* mutant. Attacker and prey bacteria were mixed in a 1:1 or 5:1 ratio and incubated for 16 hours on *Brucella* agar at 37°C under microaerobic conditons to allow competition to occur. Bacteria was harvested, serially diluted and spotted onto selection plates. The CFUs of the recovered input and output bacteria were enumerated and used to calculate the competitive index. Experiments were repeated in three biological replicates with three technical replicates. Asterisks denote a statistically significant difference (*= *p* < 0.05; ** = *p* < 0.01; *** = *p* < 0.001; **** = *p* < 0.0001).

To confirm these observations were a consequence of T6SS activity and functionality, control competition experiments were conducted with attacker 488 *gfp* and 488 *tssD* mutant or 488 *tssD+* complement as prey (Supplementary Fig. S3). In contrast there was no observed antagonism or reduction in prey survival during co-culture between kin/sister cells. This result is typical of the T6SS-mediated activity as closely related cells encode immunity proteins which protect against self-intoxication (Robitaille et al., 2021). Collectively, these findings indicate the *C. jejuni* 488 T6SS is capable of mediating inter-strain bacterial competition with susceptible T6SS-negative *C. jejuni* strains.

### *C. jejuni* 488 T6SS antagonism requires cell-to-cell contact

A cell-impermeable membrane was introduced to separate the attacker and prey cell to explore if the activity of the *C. jejuni* 488 T6SS was reliant on cell-to-cell contact. As shown in Fig. 2 and Supplementary Fig. S4, the presence of a barrier between the attacker and prey *C. jejuni* abrogated the competitive advantage and lead to increased prey survival regardless of whether the attacking strain possessed a fully functional T6SS. In fact, the presence of the membrane resulted in no statistically relevant difference between respective recovered prey input and output CFUs (Supplementary Fig. S4). Fig. S2, reveals that only during competition with the attacker 488 *tssD* mutant and the prey C*. jejuni* 81-176 and between the T6SS-negative 81-176 and 11169H *kpsM* mutant, were the differences in the competitive index comparable regardless of if the membrane was present or not. These observations indicate that the *C. jejuni* 488 T6SS mediates antagonism in a contact-dependent manner.

**Figure 2:**
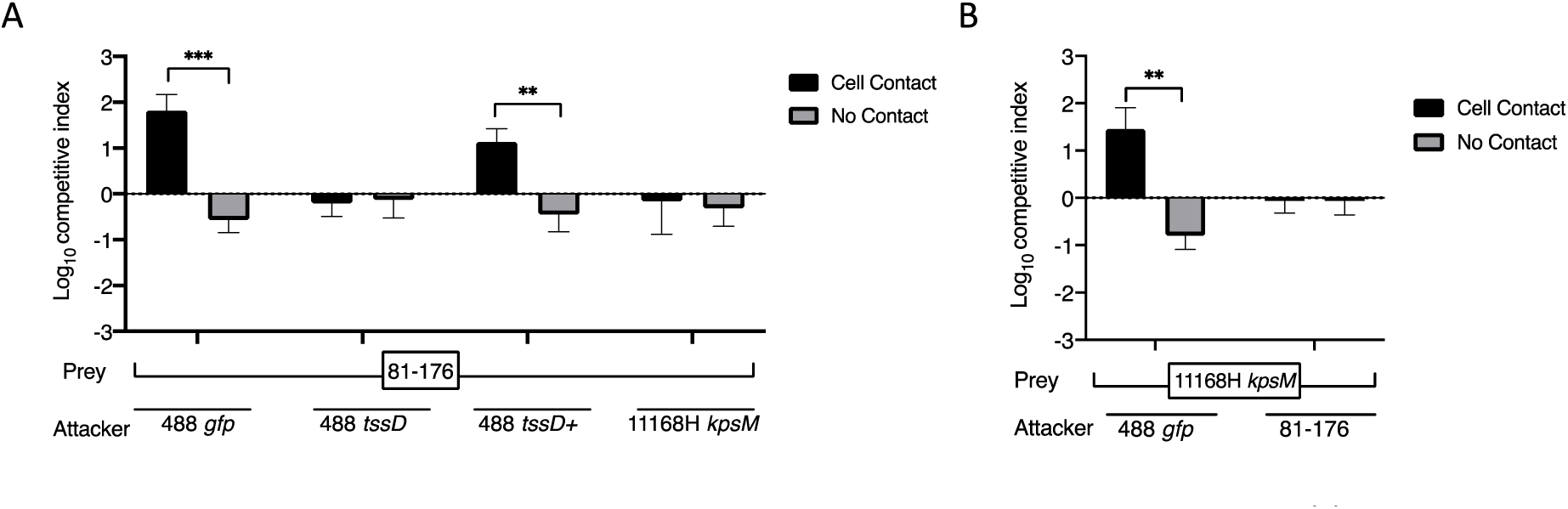
*C. jejuni* 488 T6SS-mediated antagonism is contact-dependent. Competition assays were conducted between the indicated *C. jejuni* attacker and (A) prey 81-176 strain or (B) prey 11168H *kpsM* mutant with an immepermeable cell membrane seperating the attacker and prey bacteria. After incubation for 16 hours on *Brucella* agar at 37°C under microaerobic conditons, the bacteria from the plate and membrane were harvested, serially diluted and spotted onto selection plates. The CFUs of the recovered input and output prey and attacker bacteria were enumerated and used to calculate the competitive index which were compared against a membrane-free incubation control where cell-to-cell contact was permitted. Experiments were repeated in three biological replicates with three technical replicates. Asterisks denote a statistically significant difference (** = *p* < 0.01; *** = *p* < 0.001).

### *C. jejuni* 488 T6SS antagonism is observed towards T6SS-negative *C. coli, E. coli*, and *E. faecium* strains

To follow this, the involvement of the *C. jejuni* T6SS in interspecies bacterial antagonism to ecologically relevant bacteria was investigated using T6SS-negative *C. coli* (M8 and C75) (Fig. 3a), *E. faecium* (Fig. 3b) and *E. coli* (Fig. 3c) as prey. The same observations were made as with the inter-strain competition experiments, *C. jejuni* strains possessing a functional T6SS (488 *gfp* and 488 *tssD+)* exhibited a competitive advantage with all assayed prey species. Recovered prey output CFUs were significantly decreased following incubation with T6SS functional strains, (supplementary Fig.S4). The *C. coli* M8 strain appeared to be more susceptible to T6SS attacks then *C. coli* C75, as greater competitive index values were recorded with both the 488 *gfp* and 488 *tssD*+ as attackers during competition with prey *C. coli* M8. Additionally during experimentation involving prey *E. coli,* perhaps owing to the differences in inherent doubling times between the bacterial species, a much higher attacker to prey cell density was required to observed *C. jejuni* 488 T6SS-mediated predation against the *E. coli* strain. As such at the 5:1 ratio of attacker to prey, unlike previous findings, there was no evidence of the T6SS functional 488 *gfp* and 488 *tssD*+ outcompeting the susceptible prey strain (Fig. 3c). This is corroborated by the recovered prey CFUs displayed in the supplementary Fig.S5c, which showed no statically significant decrease in recovered prey following co-incubation when compared to the unmixed input control incubated in the absence of the attacker. All together these results indicate there are inter-strain and interspecies variation in response to *C. jejuni* 488 T6SS predatory activity.

**Figure 3:**
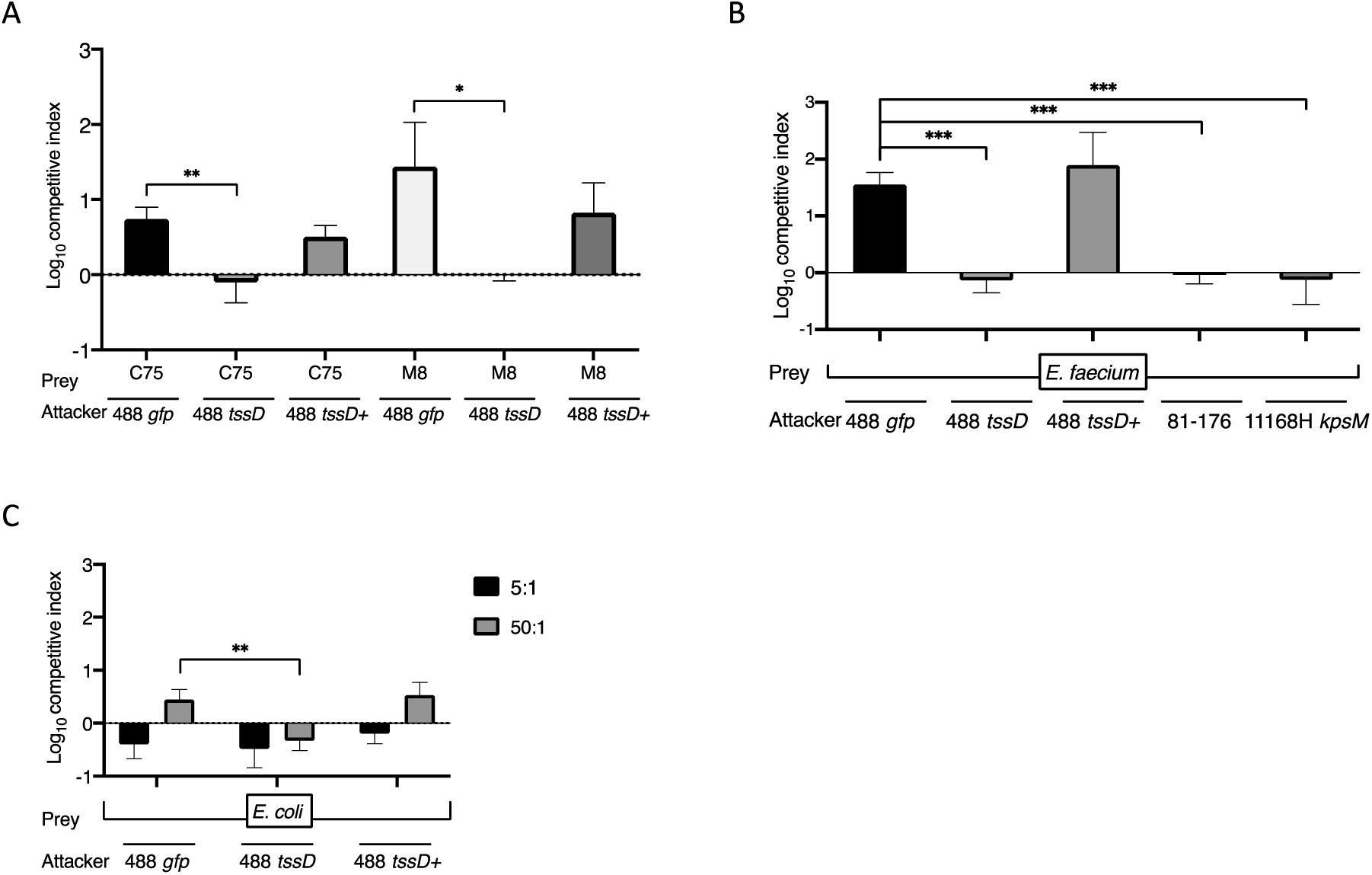
*C. jejuni* 488 T6SS functionality is required to confer a competitive advantage during competition with T6SS-negative *C. coli*, *E. faecium* and *E. coli*. Indicated attacker *C. jejuni* strains and prey (A) *C. coli* C75, *C. coli* M8 or (B) *E. faecium* were mixed in a 5:1 and incubated on 1.5% *Brucella* agar for 16 hours or (C) with prey or *E. coli* in a 50:1 ratio and incubated on Mueller Hinton agar for 16 hours. Bacteria was harvested, serially diluted, and and spotted onto selection plates. The CFUs of the recovered input and output bacteria were enumerated and used to calculate the competitive index. Experiments were repeated in three biological replicates with three technical replicates. Asterisks represent a statistically significant difference (*= *p* < 0.05; ** = *p* < 0.01; *** = *p* < 0.001)

### CJ488_0980 and CJ488_0982 modulate *C. jejuni* 488 T6SS-mediated inter-strain bacterial competition

Previous *in silico* analysis of the 488 genome CJPI-1 PAI identified CJ488_0980 and CJ488_982, which are both predicted to be putative effectors possessing domains belonging to the restriction endonuclease family Tox-REase-7 (PF15649) (Robinson et al., 2021). It was proposed that CJ488_0981 and CJ488_983 which are encoded downstream of CJ488_0980 and CJ488_982 respectively, may function as cognate immunity proteins. Prompting us to hypothesise that the CJ488_0980 and CJ488_982 putative effectors have anti-bacterial functions and are responsible for the observed *C. jejuni* 488 T6SS-mediated antagonism.

To explore if CJ488_0980 and CJ488_0982 contribute to the 488 T6SS predation activity, competition assays were performed using constructed isogenic putative effector mutants, 488 *cj0980* and 488 *cj0982*. To begin, we investigated if mutation of these putative effectors caused polar effects on T6SS functionality or increased susceptibility to kin T6SS predation. Whilst there was evidence of slight growth defects in the effector mutants with respect to the 488 wild-type strain (Supplementary Fig. S6a), there was no effect on TssD secretion, and therefore T6SS functionality through western blot analysis (Supplementary Fig. S6b). Additionally putative effector mutants were resistant to being outcompeted by the *C. jejuni* 488 T6SS, showing mutation did not impact self-intoxication protection by immunity proteins (Supplementary Fig. S7).

Notably Fig. 4a shows that loss of functionality of CJ488_0980 and CJ488_0982 led to a diminished ability of the 488 *cj0980* mutant and 488 *cj9082* mutant to outcompete the prey 81-176 strain when compared to the 488 *gfp* as an attacker. This was most pronounced during competition with the 488 *cj0980* mutant as there was almost a 2-log decrease in competitive index values recorded at the 5:1 ratio in relation to values measured with 488 *gfp* as an attacker and prey 81-176 strain. Recovered CFU counts mirror this as at the 5:1 ratio, far greater prey CFUs were recovered following co-culture with 488 *cj0980* mutant as attacker when compared to co-culture with 488 *gfp* as attacker, with almost 1-log increase in prey CFU recovery (Fig. S8). However, while similar trends were observed during competition with 488 *cj0982* as an attacker against prey 81-176, following competition with prey 11168H *kpsM* there were no statistically significant difference in the ability of the 488 *cj0982* mutant to outcompete the prey 11168H *kpsM* mutant when compared to the 488 *gfp* strain (Fig. 4b). Similarly, the recovered prey CFUs were indistinguishable following incubation with either 488 *gfp* or 488 *cj0982* as the attackers. Collectively these results suggest that whilst the CJ488_0982 contributes to the T6SS-mediated antagonism displayed by the *C. jejuni* 488 strain it does not seem to play as a critical role as the CJ488_0980 effector. However, due to the 488 *cj0980* mutant harbouring the same antibiotic selection markers as the prey 11168H *kpsM* mutant competition assays could not be performed.

**Figure 4:**
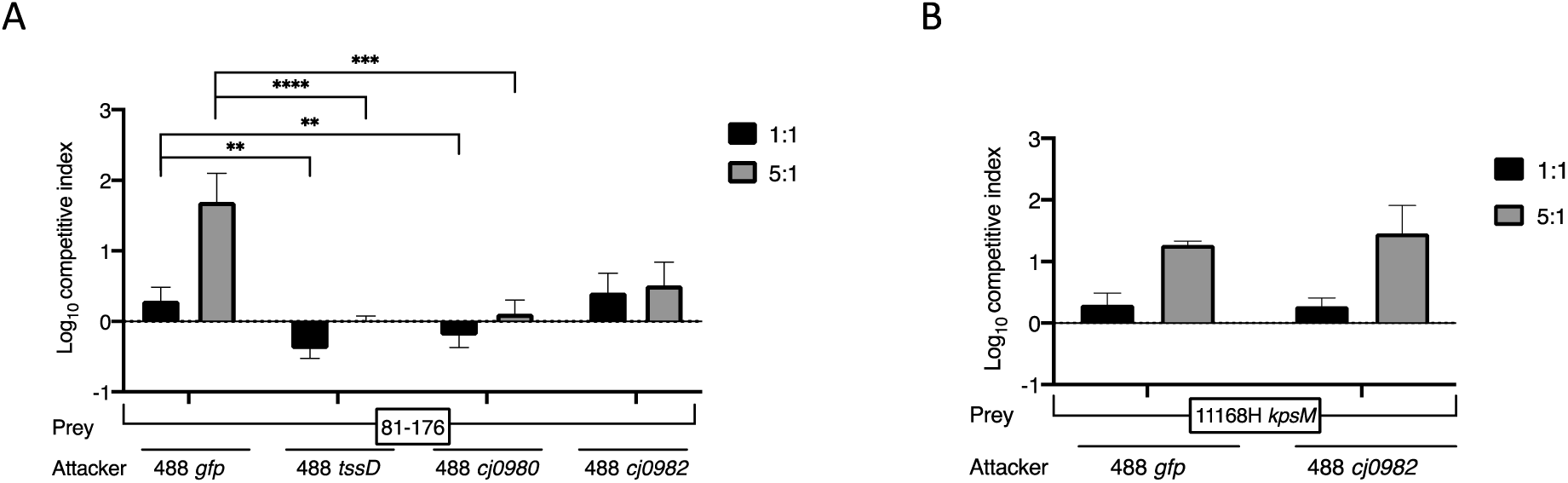
The *C.jejuni* 488 putative effectors CJ488_0980 and CJ488_0982 play a role in inter-strain T6SS-mediated attacks. Competition assays were conducted between the indicated *C. jejuni* attacker and (A) prey 81-176 strain or (B) prey 11168H *kpsM* mutant. Attacker and prey bacteria were mixed in a 1:1 or 5:1 ratio and incubated for 16 hours on *Brucella* agar at 37°C under microaerobic conditons to allow competition to occur. Harvested bacteria were serially diluted and spotted onto selection plates. The CFUs of the recovered attacker and prey input and output were enumerated and used to calculate the competitive index values. Experiments were repeated in three biological replicates with three technical replicates. Asterisks denote a statistically significant difference (** = *p* < 0.01; *** = *p* < 0.001; **** = *p* < 0.0001).

### CJ488_0980 and CJ488_0982 functionality is required for *C. jejuni* 488 T6SS-mediated interspecies bacterial competition

Equally, the 488 *cj0980* mutant and 488 *cj0982* mutant displayed reduced ability to outcompete prey *C. coli* and *E. faecium* when compared to the 488 *gfp* attacker (Fig. 5a and Fig. 5b). Reduced competitive advantage against the *C. coli* and *E. faecium* strains by the putative effector mutants are not to the degree of 488 *tssD* mutant. This indicates that while loss of functionality of CJ488_0980 and CJ488_0982 reduced’ the ability of the T6SS to outcompete competitors it does not completely abrogate T6SS-mediated antagonism. Therefore, there are likely other effectors being utilised.

**Figure 5:**
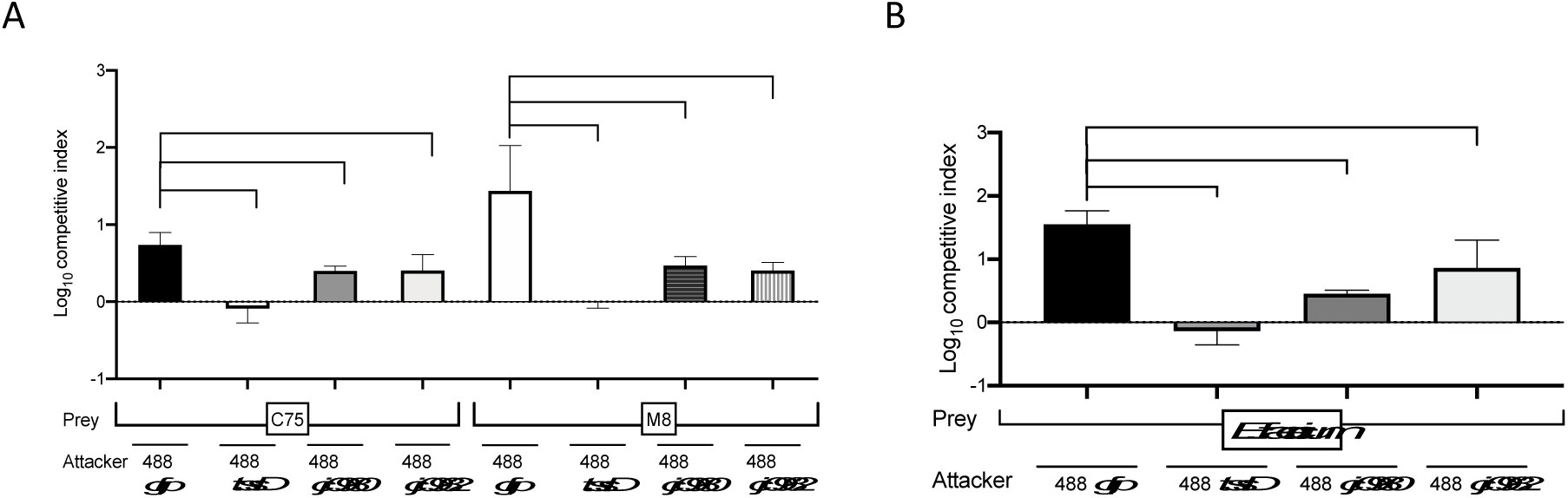
The *C.jejuni* 488 putative effectors CJ488_0980 and CJ488_0982 mediate interspecies T6SS-mediated attacks. Competition assays were conducted between the indicated *C. jejuni* attacker and (A) prey *C. coli* C75 or *C. coli* M8 or (B) *E. faecium*. Attacker and prey bacteria were mixed in a 5:1 ratio and incubated for 16 hours on *Brucella* agar at 37°C under microaerobic conditons to allow competition to occur. Bacteria were harvested, serially diluted and spotted onto selection plates. The CFUs of the output recovered attacker and prey input and output bacteria were enumerated and used to calculate the competitive index values. Experiments were repeated in three biological replicates and three technical replicates. Asterisks denote a statistically significant difference (\**p* = < 0.05; ** = *p* < 0.01; *** = *p* < 0.001).

### The *C. jejuni* 488 T6SS modulates host cell interactions during infection of Human Intestinal Epithelial cells

Previously the *C. jejuni* 488 T6SS was revealed to have a role in enhancing interaction with primary chicken cells as well as potential roles in pathogenesis and direct cytotoxicity to equine erythrocytes (Liaw et al., 2019, Bleumink-Pluym et al., 2013). To gain further insight in T6SS responses to human IECs we examined the expression of *tssB*, *tssC* and *tssD* after infection of T84 and Caco-2 cells with 488 wild-type strain (Supplementary 9). qRT-PCR results revealed all three genes were upregulated in the presence of both cell types 3 hours post-infection when compared to the cell monolayer-free control.

To further investigate whether the activity of a functional T6SS modulates the ability of 488 strain to interact (adhere and invade) with human IECs, infection assays were conducted. At 3 hours post infection, significant decreases in host cell interaction in both intestinal cell lines were observed during infection with the 488 *tssD* mutant when compared to infection with the 488 wild-type strain (Fig. 6a and Fig.6b). Conversely, these reductions in interactions were not observed following infection with the 488 *tssD*+ complement, which interacted with both cell types at similar levels as 488 wild-type strain. Similarly, Fig.6c showed there were marked decreases in invasive capacity recorded with the 488 *tssD* mutant with respect to the wild-type strain in the T84 cell line. Of note this phenotype was absent during infection in the Caco-2 cell line, the 488 wild-type strain, 488 *tssD* mutant and 488 *tssD*+ complement displayed no statistically significant differences invasive capacity to Caco-2 cells (Fig.6d). Lastly, intracellularly survival assays revealed an interesting phenotype, inversely the 488 *tssD* mutant displayed an enhanced ability to survival intracellularly when compared to both the 488 wild-type strain and 488 *tssD+* complement (Fig. 7). These results indicate that whilst the 488 T6SS appears to modulate the ability of the strain to interact with and invade human IECs, the T6SS may play a unique role in intracellular interactions and long-term survival.

**Figure 6:**
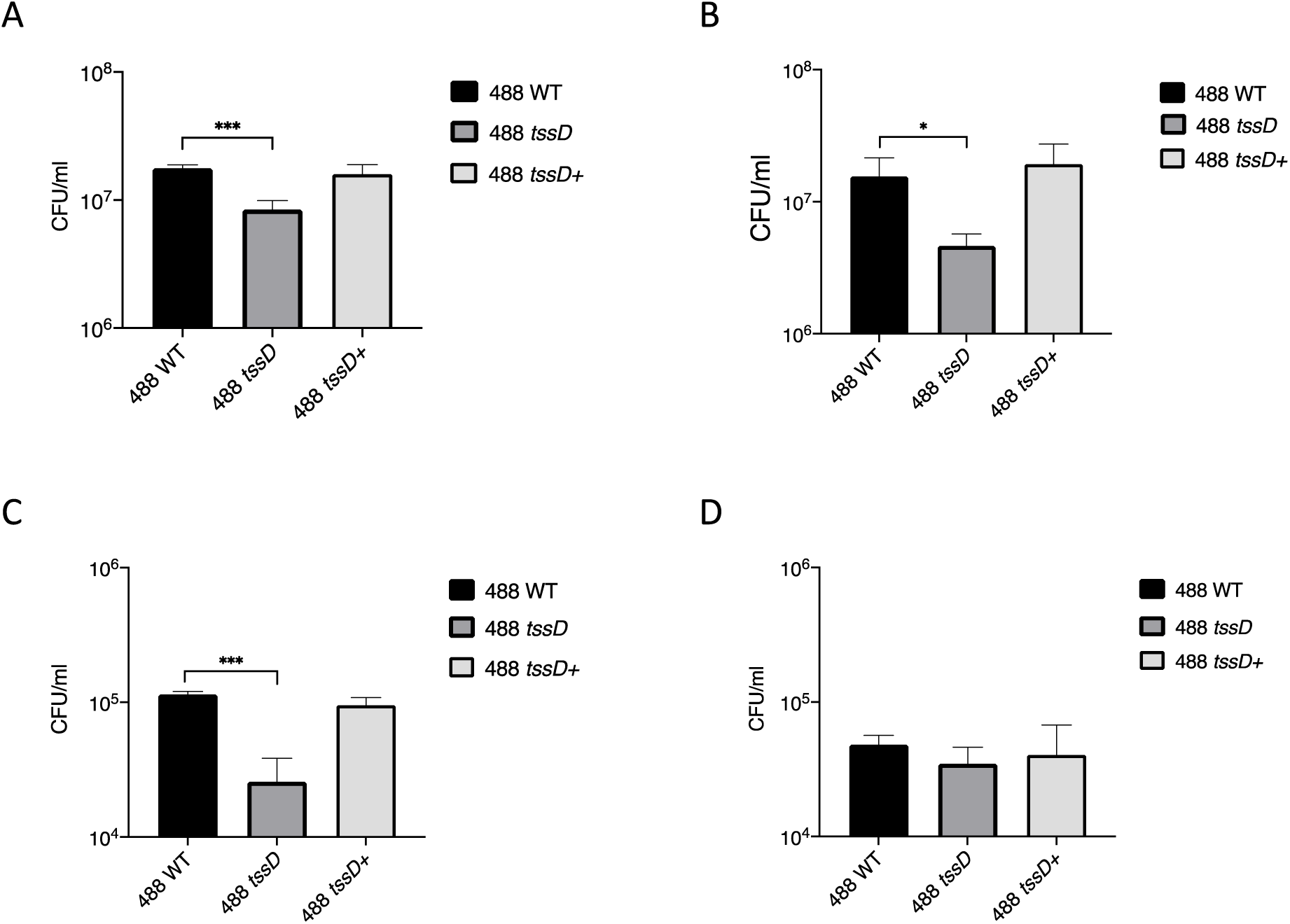
Interactions with and invasion of T84 and Caco-2 intestinal epithelial cells by *C. jejuni* 488 wild-type strain, 488 *tssD* mutant and 488 *tssD+* complement. (A, C) T84 and (B, D) Caco-2 cells were infected with *C. jejuni* 488 wild-type strain, 488 *tssD* mutant or 488 *tssD+* complement (MOI 200:1) for 3 hours at 37°C in a 5% CO_2_ incubator. (A, B) For interaction assays the cells were then washed with PBS, lysed with 0.1% (v/v) Triton X-100. (C, D) For invasion assays, after the initial 3 hour infection the cell monolayers were incubated with gentamicin (150 μg/ml) for 2 hours to kill extracellular bacteria and then lysed with 0.1% (v/v) Triton X-100. For both interaction and invasion assays the cell lysates were serially diluted and then CFU/ml were counted. Experiments were repeated in three biological replicates with three technical replicates. Asterisks denote a statistically significant difference (*** = *p* < 0.001).

**Figure 7:**
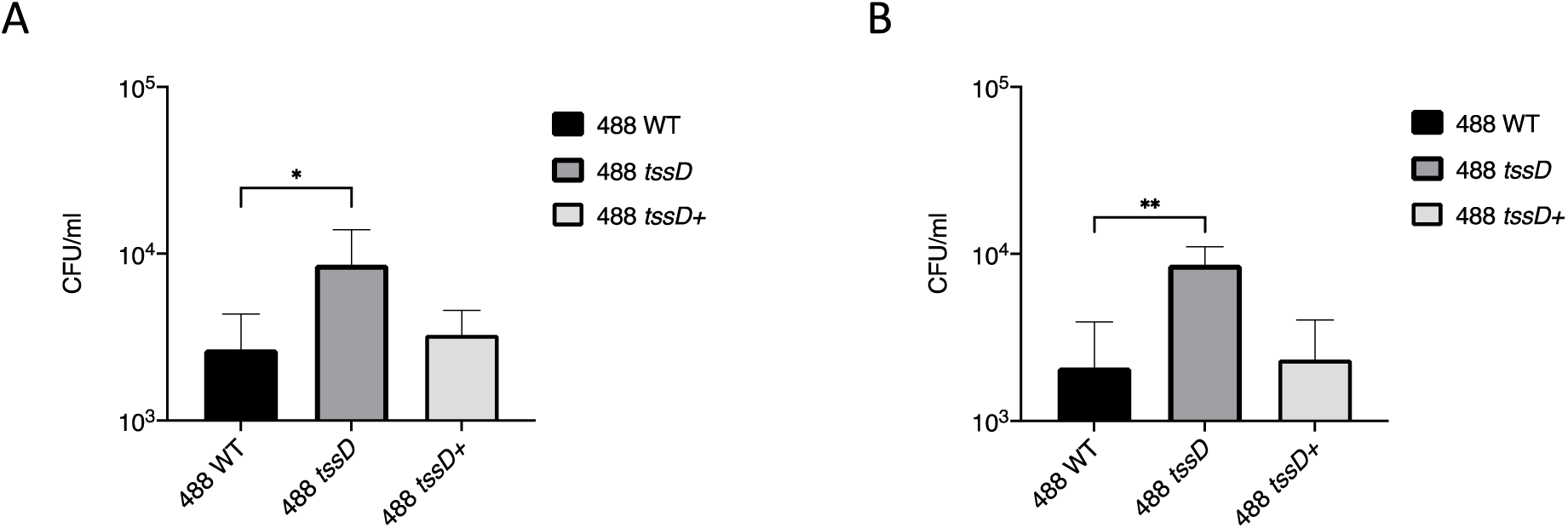
Intracellular survival of *C. jejuni* 488 wild-type strain, 488 *tssD* mutant and 488 *tssD+* complement during infection of T84 and Caco-2 intestinal epithelial cells. (A) T84 or (B) Caco-2 cells were infected with *C. jejuni* 488 wild-type strain, 488 *tssD* mutant or 488 *tssD+* complement (MOI 200:1) for 3 hours at 37°C in a 5% CO_2_ incubator. The cell monolayers were incubated with gentamicin (150 μg/ml) for 2 hours to kill extracellular bacteria and then further incubated with gentamicin (10 μg/ml) for 18 hours. The T84 and Caco-2 cells were then lysed with 0.1% (v/v) Triton X-100, cell lysates were serially diluted and then CFU/ml were counted. Experiments were repeated in three biological replicates with three technical replicates. Asterisks denote a statistically significant difference (*= *p* < 0.05; ** = *p* < 0.01).

## Discussion

Research on the *C. jejuni* T6SS has lagged other enteric bacteria, with much of the investigations focussing on prevalence and the only effector characterised experimentally so far being TssD (Robinson et al., 2021). Despite several *in silico* bioinformatics analyses have identified candidate putative effector proteins through *in silico* analysis, no *C. jejuni* T6SS effectors have been experimentally investigated (Robinson et al., 2021, Katz et al., 2023, Robinson et al., 2022). The T6SS has been demonstrated to contribute to the pathogenic potential of T6SS-positive isolates the presence of The T6SS is associated with phenotypes of enhanced virulence in the host and survival during meat processing and storage (Marasini et al., 2020, Bleumink-Pluym et al., 2013). However, as the functions of the T6SS are inextricably linked to the repertoire of effectors that are encoded and secreted by the bacteria, it is hoped that probing of the T6SS effectors may further elucidate the role(s) of the *C. jejuni* secretion system.

Whilst seminal studies initially ascribed the T6SS to a function of aiding pathogenesis, it has become more widely accepted that the primary role of most encoded T6SS is to modulate microbial interactions and more specifically mediate antagonism towards competitors (Coulthurst, 2019, Drebes Dorr and Blokesch, 2018). Recent studies have confirmed that *C. jejuni* strains isolated from broiler chickens’ caecal content predate on *E. coli* DH5, although T6SS-positive strains incur a predation cost under bile stress through interbacterial competition assays (Gupta et al., 2021, Gupta et al., 2024). Our findings corroborate this as the *C. jejuni* 488 strain displayed T6SS-mediated antagonistic behaviour not only to *E. coli*, but to more closely related *C. coli* strains and Gram-positive *E. faecium*. These bacteria are all ecologically relevant as they can be found as competitors in the same diverse polymicrobial niches that *C. jejuni* reside such as the intestinal tract of animals and the environment (Rivera-Gomis et al., 2021, Lu et al., 2003). T6SS-mediated activity against the Gram-positive *E. faecium* strain confirms a more widely agreed consensus that T6SS-mediated killing can be targeted towards Gram-positive bacteria (Hersch et al., 2020). The *A. baumanii* T6SS has been shown to deploy attacks against Gram-positive *Bacillus subtilis*, *Listeria monocytogenes* and *Staphylococcus aureus* using the Tse4 effector, which has both lytic transglycosylase and endopeptidase activities (Le et al., 2021). This finding widens the conceived limits of the influence of the *C. jejuni* T6SS may have prokaryotic interactions in polymicrobial communities and so warrants further investigation.

Data presented in this study demonstrates the first example of the *C. jejuni* T6SS being utilised for inter-strain competition as the *C. jejuni* 488 strain is capable of outcompeting T6SS-negative strains 11168H *kpsM* and 81-176. Examples of T6SS-mediated territorial behaviour between strains of the same species have been observed previously, most famously with *Proteus mirabilis* which deploys the T6SS for kin-recognition which results in the formation of distinct Dienes lines between competing *P. mirabilis* strains (Wenren et al., 2013, Alteri et al., 2013). The T6SS can distinguish between self and non-self due to the presence and/or absence of effectors and their cognate immunity proteins. This has implications on population dynamics for both T6SS-negative and non-kin T6SS-positive isolates (Kostiuk et al., 2021). T6SS duelling may even occur between T6SS-positive non-kin strains that possess different effector and immunity protein repertoires; future studies can investigate competition between T6SS-positive *C. jejuni* strains (Kostiuk et al., 2021).

Attacking strains required not only a fully functional T6SS to facilitate predation, *C. jejuni* 488 T6SS-mediated attacks were also revealed to be dependent on cell-to-cell contact. The physical separation of the attacker and prey through a cell-impermeable membrane abolished predation. However, this does not necessarily mean the *C. jejuni* T6SS activity is exclusively reliant on cell-contact. Whilst cell contact was originally thought to be a fundamental characteristic of T6SS-mediated attacks, it has been revealed that T6SS may function in a contact-independent manner (Song et al., 2021, Si et al., 2017). Contact-independent roles of the T6SS are typically associated with metal acquisition from the extracellular milieu but can be linked to indirect exploitative competition (Yang et al., 2021). However, recently the *Yersina pseudotuberculosis* effector CccR has been found to mediate direct interbacterial competition in a contact-dependent and contact-independent manner (Wang et al., 2022).

The *C. jejuni* 488 T6SS was additionally shown display antagonistic behaviour in a density-dependent manner towards susceptible prey. This is consistent with other bacterial T6SS as a higher cell density allows increased and earlier cell contact (Smith et al., 2020). This was demonstrated when a substantially higher attacker to prey ratio of 50:1 was required to elicit *C. jejuni* T6SS-mediated competition with *E. coli* compared to the other bacterial species. The 50:1 ratio may also be reflective of the incompatibility between the growth rates of *C. jejuni* and *E. coli.* Fast-growing *E. coli* microcolonies can grow at a rate that outpaces T6SS intoxication, and this has been shown to be an effective T6SS defence strategy against T6SS attacks by *Vibrio cholerae* (Borenstein et al., 2015).

As of yet none of the reported phenotypes of *C. jejuni* T6SS have been associated with any specific cargo effectors. As such this characterisation of the putative effectors CJ488_980 and CJ488_0982 represents the first example of a “classical” cargo effector characterisation in *C. jejuni.* Loss of function of CJ488_980, CJ488_0982, diminished but did not completely abolish the competitive advantage measured during co-culture with T6SS-negative *C. jejuni*, *C. coli* and *E. faecium*, unlike competition with the non-functional T6SS 488 *tssD* mutant. Therefore, we infer CJ488_0980 and CJ488_0980 contributes to the 488 strain T6SS killing of competitors but are not solely responsible. Greater losses in competitive advantage against prey 81-176 were observed with the 488 *cj0980* mutant than the 488 *cj0982* mutant, so we propose these effectors have a synergistic effect, each contributing to bacteria antagonism but CJ488_0980 may play a more critical role.

CJ488_0980 and CJ488_0980 are predicted to possess Tox-REase-7 domains with a conserved restriction endonuclease fold. No T6SS effectors with this characteristic fold have currently been characterised. However, examples of Tox-REase-7-fold domain containing proteins include the CdiA identified in *Pseudomonas* spp. and *Acinetobacter* spp. which are linked to the activity of T5SS contact-dependent growth inhibition (CDI) systems (De Gregorio et al., 2019, Mercy et al., 2016). The *C*-terminal of CdiA2 of the *Acinetobacter baumanii* DSM30011 strain displays nuclease activity by causing multiple DNA breaks to the chromosomal DNA, arrestment of cell division, inhibition of growth and ultimately cell death when expressed in *E. coli* (Roussin et al., 2019). Due to the shared restriction endonuclease domain, we hypothesise the CJ488_0980 and CJ488_0980 may elicit similar responses and target the DNA or RNA of recipient prey cells, but further characterisation is needed to confirm this.

Interestingly a study looking at molecular epidemiology and the prevalence of putative effectors amongst Chilean clinical strains found CJ488_0980 and CJ488_0982 were not highly conserved (Katz et al., 2023). Using hierarchical clustering analysis based on repertoire of putative effector proteins and immunity five major subgroups (I to V) were formed, while the presence of CJ488_0982 was restricted was subgroups II and III, CJ488_0980 was not presence in any of the Chilean isolates (Katz et al., 2023). This is evidence of the diversity of *C. jejuni* effector repertoires and the potential implications this has on microbial dynamics and the virulence potential of the strains. Further work needs to be undertaken to further probe the mode of action of this putative effector as well as to characterise other *C. jejuni* T6SS effectors to grasp how they contribute to overall virulence of the isolates that harbour them.

Data presented in this study indicates that *C. jejuni* 488 T6SS functionality correlates with enhanced ability to interact and invade human IECs. This supports work conducted using *in vitro* primary chicken cell lines and *in vivo* infections of broiler chickens and is consistent with infection of RAW 267.4 macrophage cell lines (Liaw et al., 2019, Lertpiriyapong et al., 2012). Increased T6SS-mediated invasive abilities were restricted to the infection of the T84 cell line, and not observed during infection of Caco-2 cells. This is likely due to cell lines variations. Although T84 and Caco-2 cells are both colonic adenocarcinoma-derived cell lines, they possess distinct morphologies, biochemistry and functionalities with T84 resembling colonocytes whereas the Caco-2 cells are more similar to enterocytes (Devriese et al., 2017).

Interestingly a phenotype of increased intracellular survival of the non-functional *tssD* mutant with respect to the 488 wild-type strain was observed in both cell types. This phenotype has yet to be described in any T6SS-positive *C. jejuni* strains but has been documented in *S. typhimurium* 14028s strain which encodes 3 TssD (Hcp) family proteins each believed to have different roles (Wang et al., 2020). The authors proposed the reduced metabolic burden on the Hcp1 mutant strain permitted increased intracellular survival in a *Dictyostelium discoideum* model (Wang et al., 2020). Similarly, *H. hepaticus* T6SS mutants displayed an increased intracellular presence and adherence to MODE-K cells (murine IEC) and *in vivo* colonisation of murine intestine, with the bacteria hypothesised to use the T6SS to limit colonisation and intestinal inflammation to promote long-term non-pathogenic symbiosis in the host (Chow and Mazmanian, 2010).Lertpiriyapong et al., have proposed that the *C. jejuni* TssD functions as an evolved effector to facilitate host cell invasion as overexpression of TssD enhanced cell adhesion and invasion in a dose-dependent manner (Lertpiriyapong et al., 2012). For *C. jejuni* the mechanisms facilitating these observed host cell modulation phenotypes have yet to be determined.

Our observations linked T6SS-mediated prey killing to the functionality of CJ488_980 and CJ488_0982 two Tox-REase-7 domain containing putative effectors. Additionally, the *C. jejuni* 488 T6SS was shown to be upregulated in the presence of T84 and Caco-2 cells, with infection assays demonstrating the *C. jejuni* 488 T6SS enhances host cell interaction, invasion and modulates intracellular survival. These initial analyses of CJ488_980 and CJ488_0982 through the constructed isogenic mutants, 488 *cj0980 and* 488 *cj0982* are the first *in vitro* characterisation of cargo effectors in a *Campylobacter* spp. Further study of these putative effectors will be crucial for providing greater insights of the role(s) of the *C. jejuni* T6SS. In summary, we have demonstrated the *C. jejuni* 488 T6SS mediates host cell dynamics and predation of non-kin strains, which has implications on virulence and interbacterial competition in the host and environment.

## Supplementary information

**Figure S1:**
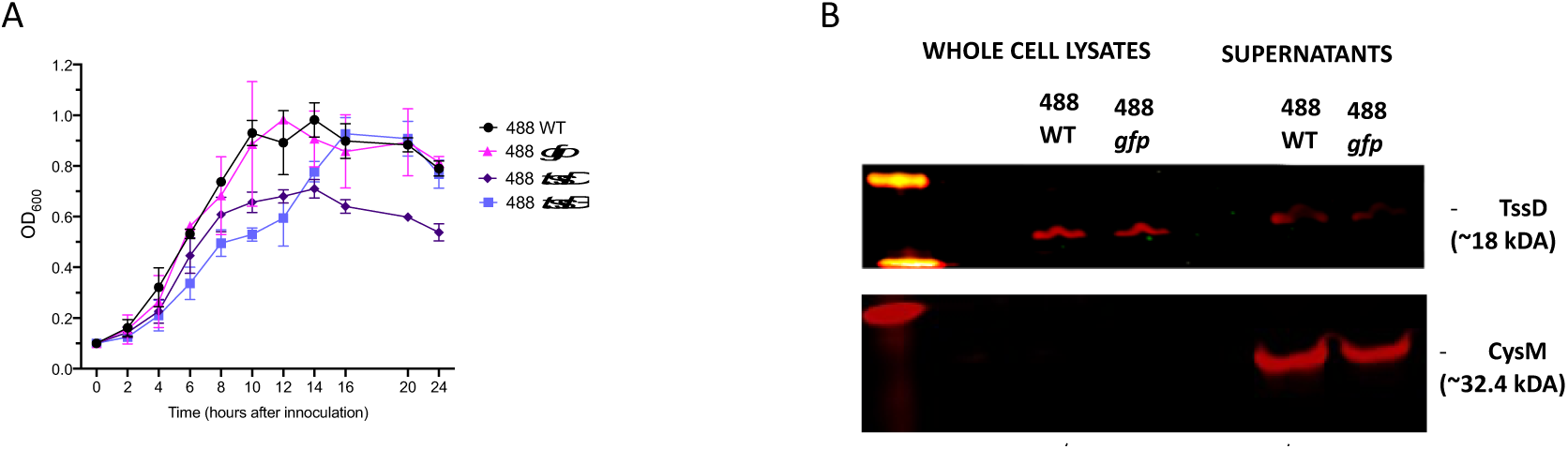
(A) OD_600_ growth kinetics of 488 wild-type strain, 488 *gfp,* 488 *tssD* mutant and 488 *tssD* + complement. *C. jejuni* was used to inoculate *Brucella* broth flasks to an OD_600_ of 0.1 and then incubated with shaking at 75 rpm at 37°C under microaerobic conditions for 24 hours. The OD_600_ of the flasks were recorded at 2,4,6,8,10,12,14,16, 20 and 24 hours after inoculation. (B) Western blot probing for secretion TssD by 488 wild-type strain (WT) and 488 *gfp*. Whole cell lysates and supernatant were prepared from *C. jejuni* 488 wild-type strain and 488 *gfp* grown in *Brucella* broth flasks for 16 hours at 37°C under microaerobic conditions whilst shaking at 75 rpm. TssD has an estimated molecular weight of 18 kDa and the lysis control CysM protein has a molecular weight of 32.4 kDa

**Figure S2:**
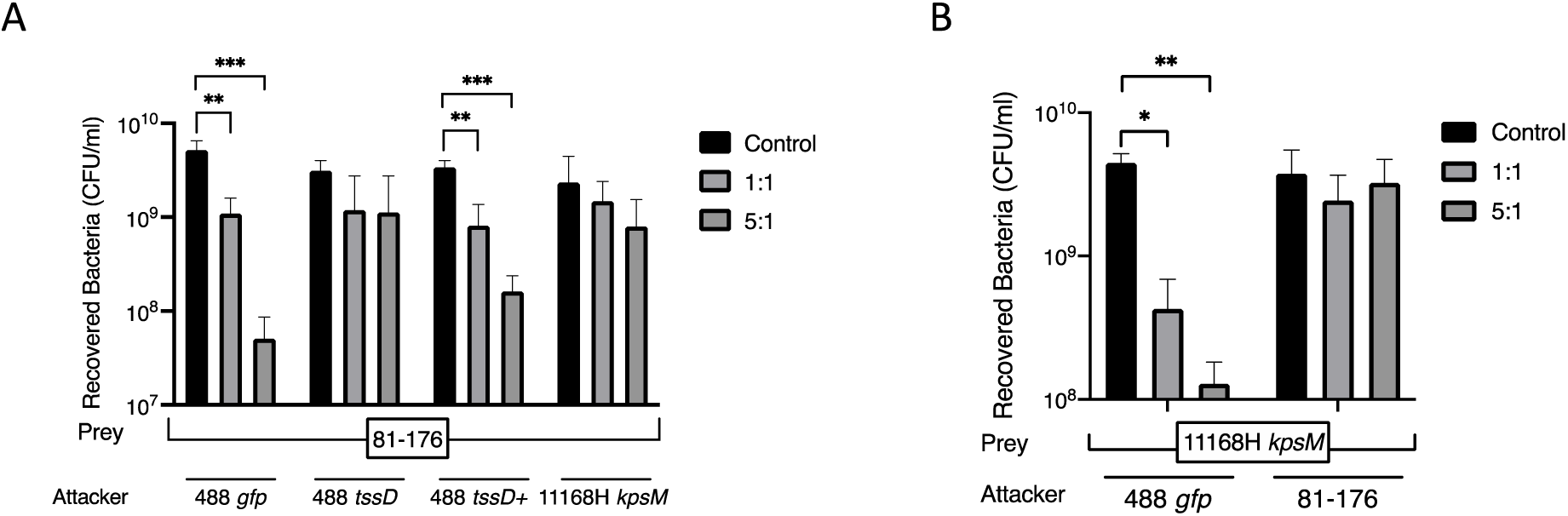
Recovered input and output prey CFU/ml following inter-strain competition assays. (A) recovered prey 81-176 CFU/ml after co-culture with 488 *gfp*, 488 *tssD* mutant, 488 *tssD+* or 111668H *kpsM* mutant as attackers in a 1:1 or 5:1 ratio of attacker to prey. (B) Recovered prey 111668H *kpsM* mutant CFU/ml after co-culture with 488 *gfp* or 81-176 strain as attackers in a 1:1 or 5:1 ratio of attacker to prey. Bacteria was co-incubated on *Brucella* agar for 16 hours at 37°C under microaerobic conditons. After incubation bacteria were harvested, serially diluted and plated onto selective agar to select for either the attacker or prey bacteria CFUs. The control represents the unmixed input of the respective prey bacteria. Experiments were repeated in three biological replicates with three technical replicates. Asterisks denote a statistically significant difference (*= *p* < 0.05; ** = *p* < 0.01; *** = *p* < 0.001).

**Figure S3:**
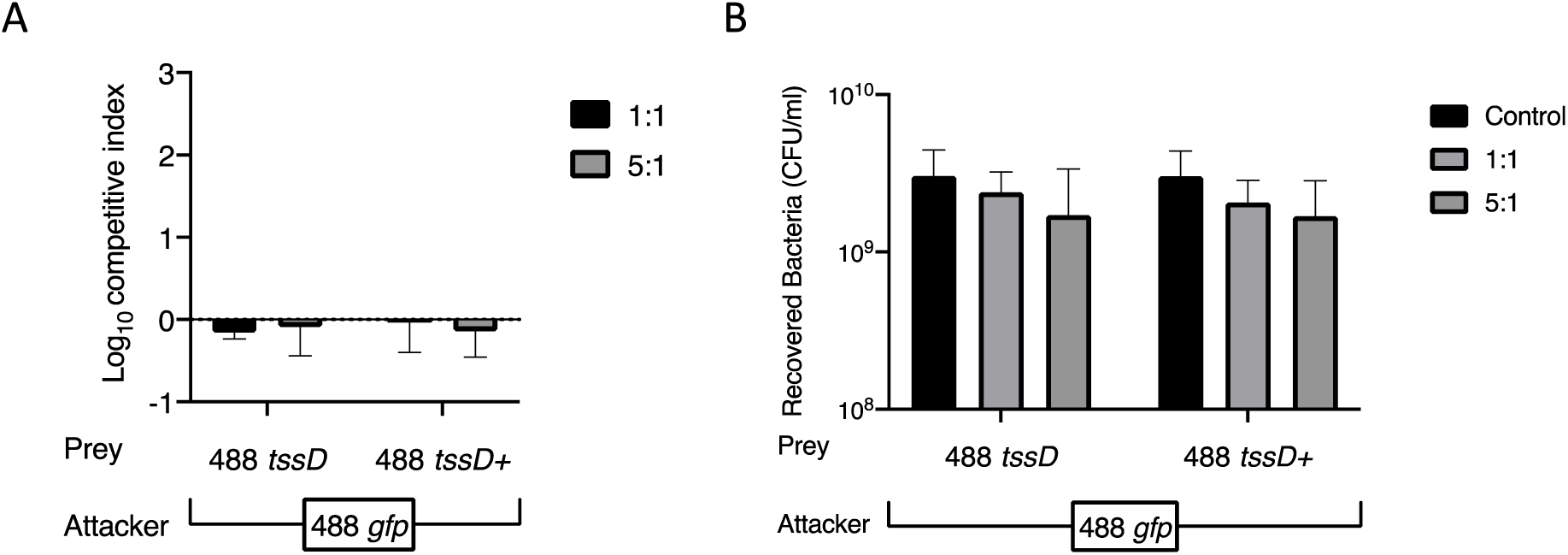
Self-intoxication control experiment for inter-strain competition assays and the associated recovered prey CFU/ml. Attacker 488 *gfp* was mixed prey 488 *tssD* mutant or prey 488 *tssD+* complement respectively in a 1:1 or 5:1 and co-incubated on *Brucella* agar for 16 hours at 37°C under microaerobic conditons. After incubation bacteria were harvested, serially diluted, and plated onto selective agar to select for either the attacker or prey bacteria. (B) Competitive index values were calculated using the recovered output and input CFUs from the attacker and (B) prey bacteria. (B) The control represents the unmixed input of the respective prey bacteria. Experiments were repeated in three biological replicates with three technical replicates.

**Figure S4:**
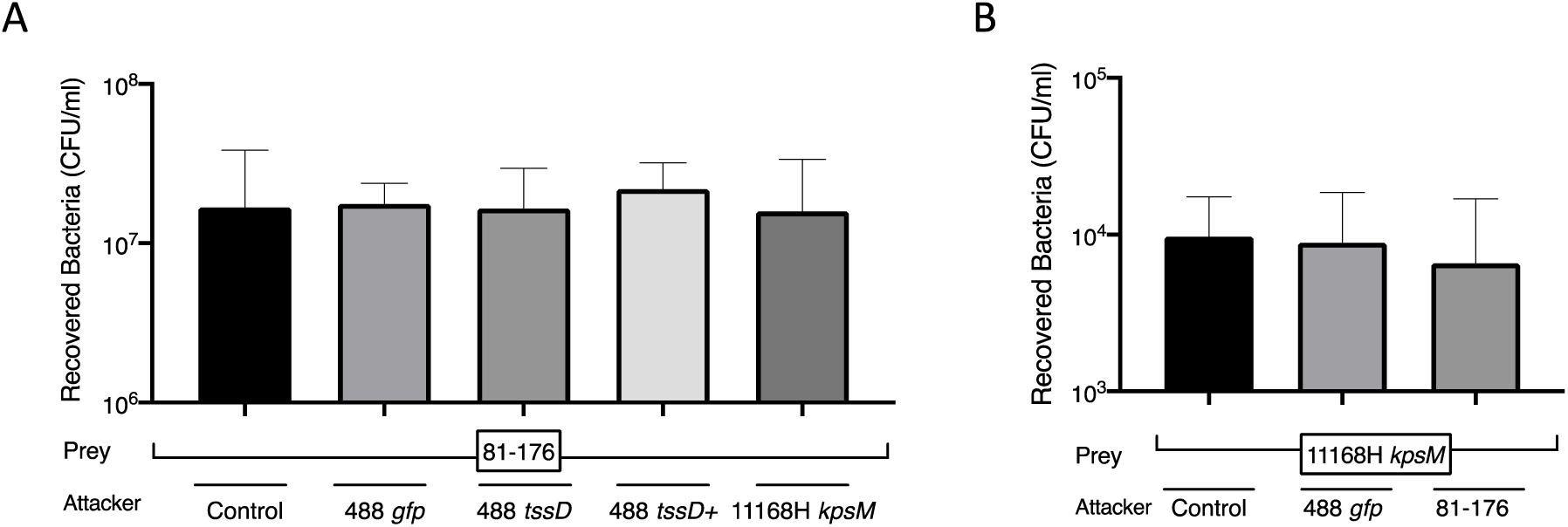
Recovered input and output prey CFU/ml following competition assays performed in the presence of a cell impermeable membrane separating the attacker and prey bacteria. (A) recovered prey 81-176 strain CFU/ml after incubation with 488 *gfp*, 488 *tssD* mutant, 488 *tssD+* complement or 11168H *kpsM* mutant as attackers in a 5:1 ratio of attacker to prey on *Brucella* agar for 16 hours at 37°C under microaerobic conditons. (B) Recovered prey 111668H *kpsM* mutant CFU/ml after incubation with 488 *gfp* or 81-176 strain as attackers in a 5:1 ratio of attacker to prey. After incubation the bacteria was harvested, serially diluted, and plated onto selective agar to select for either the attacker or prey bacteria The control represents the unmixed input of the respective prey bacteria. Experiments were repeated in three biological replicates with three technical replicates.

**Figure S5:**
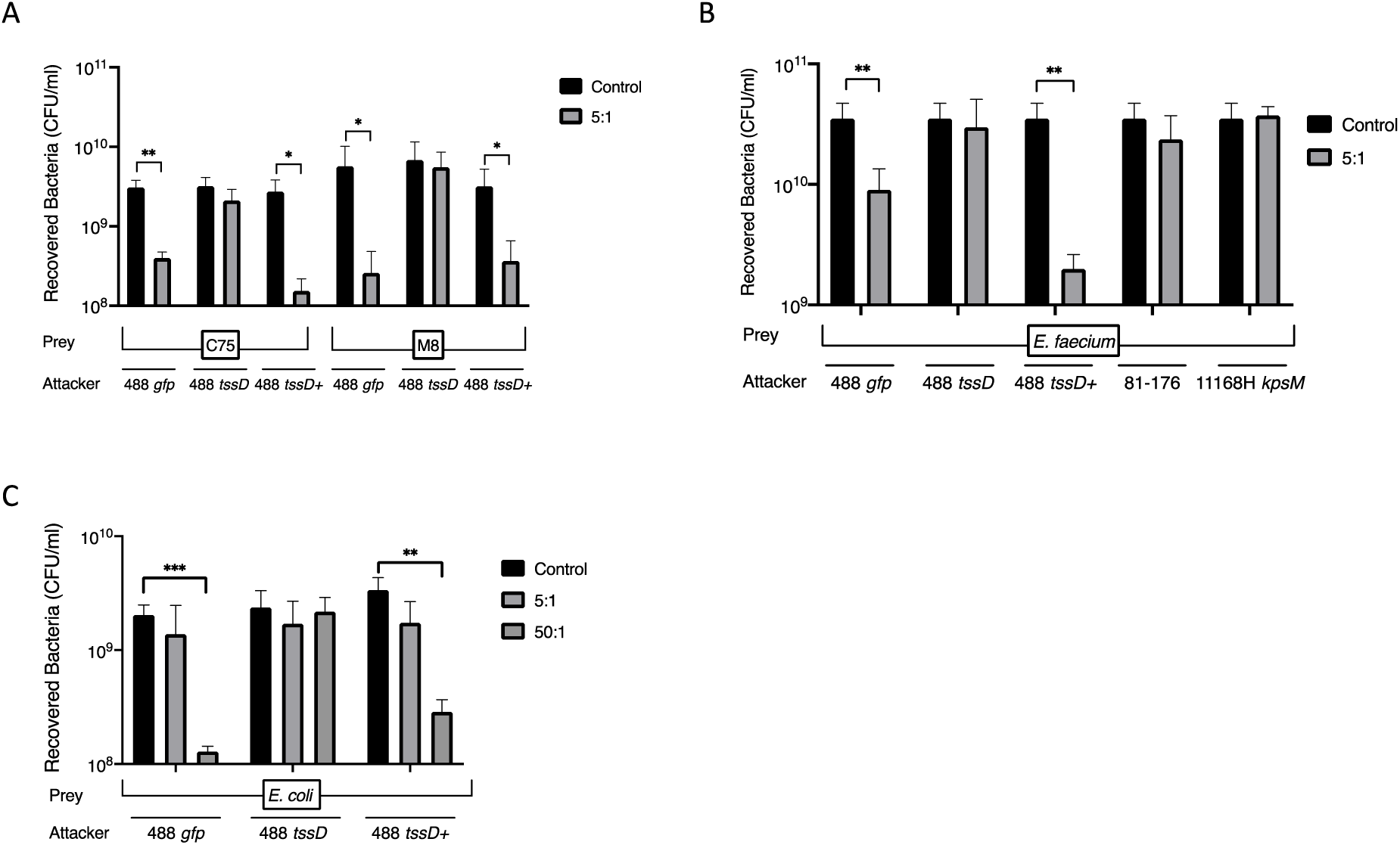
Recovered input and output prey CFU/ml following interspecies competition assays. The 488 *gfp, tssD* mutant or *tssD+* complement respectively as attackers were mixed and incubated with prey in a 5:1 ratio and incubated on *Brucella* agar for 16 hours at 37°C under microaerobic conditons or (C) 50:1 ratio of attacker to prey and co-incubated on Mueller Hinton agar for 16 hours at 37°C under microaerobic conditons. After incubation bacteria were harvested, serially diluted, and plated onto selective agar to select for either the attacker or prey bacteria. The recovered input and output CFU/ml of the prey (A) *C. coli* C75 and *C. coli* M8 strains, (B) *E. faecium* or (C) *E. coli* are displayed and compared against the respective unmixed input control incubated in the absence of the attacker. Experiments were repeated in three biological replicates with three technical replicates. Asterisks denote a statistically significant difference (*= *p* < 0.05; ** = *p* < 0.01; *** = *p* < 0.001).

**Figure S6:**
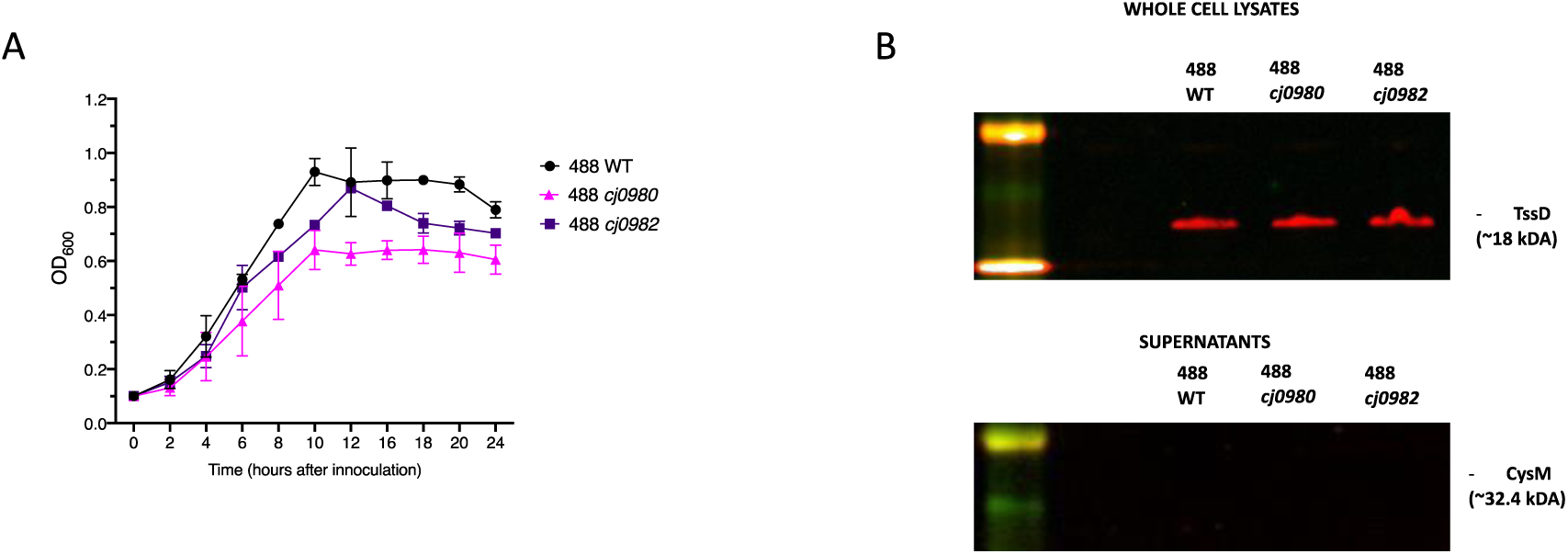
(A) OD_600_ growth kinetics of *C. jejuni* 488 wild-type strain, 488 *cj0980* mutant and 488 *cj0982* mutant*. C. jejuni* was used to inoculate *Brucella* broth flasks to an OD_600_ of 0.1 and then incubated with shaking at 75 rpm at 37°C under microaerobic conditions for 24 hours. The OD_600_ of the flasks were recorded at 2,4,6,8,10,12,14,16, 20 and 24 hours after inoculation.(B) Western blot probing for TssD secretion by *C. jejuni* 488 wild-type strain (WT), 488 *cj0980* mutant and *cj0982* mutant. Whole cell lysates and supernatant were prepared from *C. jejuni* 488 wild-type strain, 488 *cj0980* mutant and *cj0982* mutant grown in *Brucella* broth flasks for 16 hours at 37°C under microaerobic conditions whilst shaking at 75 rpm. TssD has an estimated molecular weight of 18 kDa and the lysis control CysM protein has a molecular weight of 32.4 kDa.

**Figure S7:**
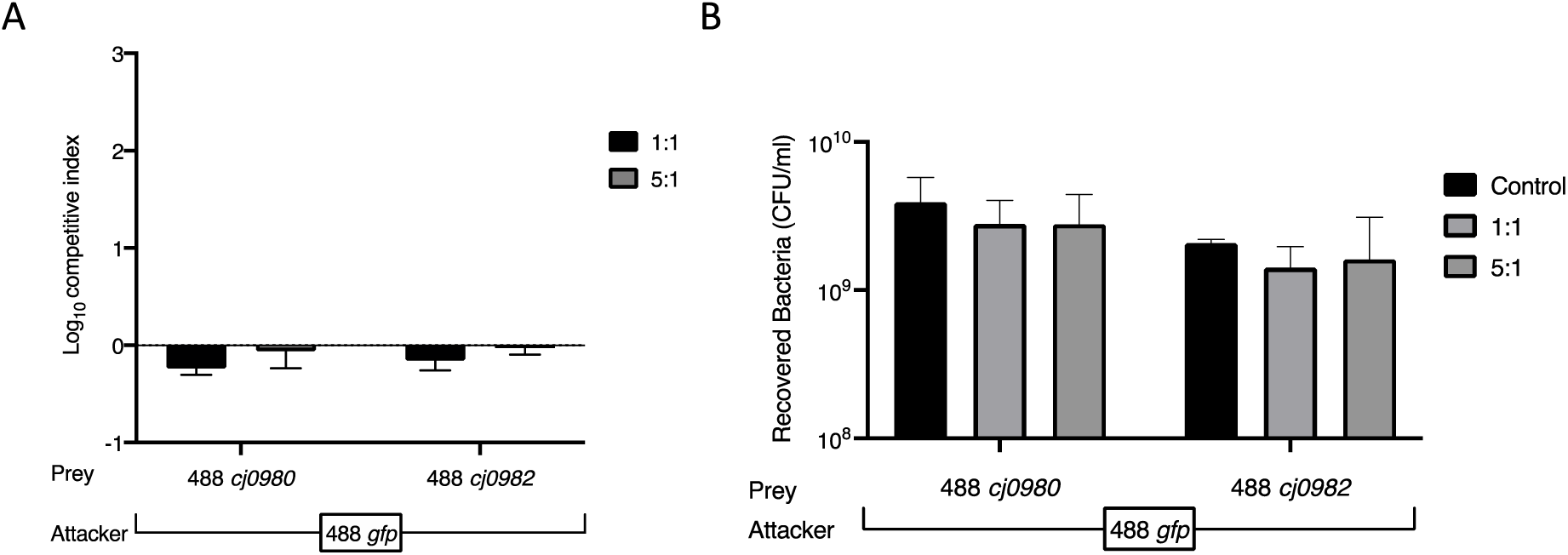
Self-intoxication control experiment with the putative effector mutants and the associated recovered prey CFU/ml. Attacker 488 *gfp* was mixed prey 488 *cj0980* mutant or 488 *cj0982* mutant in a 1:1 and 5:1 and co-incubated on *Brucella* agar for 16 hours under microaerobic conditions. After incubation bacteria were harvested, serially diluted, and plated onto selective agar to select for either the attacker or prey bacteria. (A) The competitive index values were calculated using the (B) recovered output and input prey CFUs. (B) The control represents the unmixed input of the respective prey bacteria. Experiments were repeated in three biological replicates with three technical replicates.

**Figure S8:**
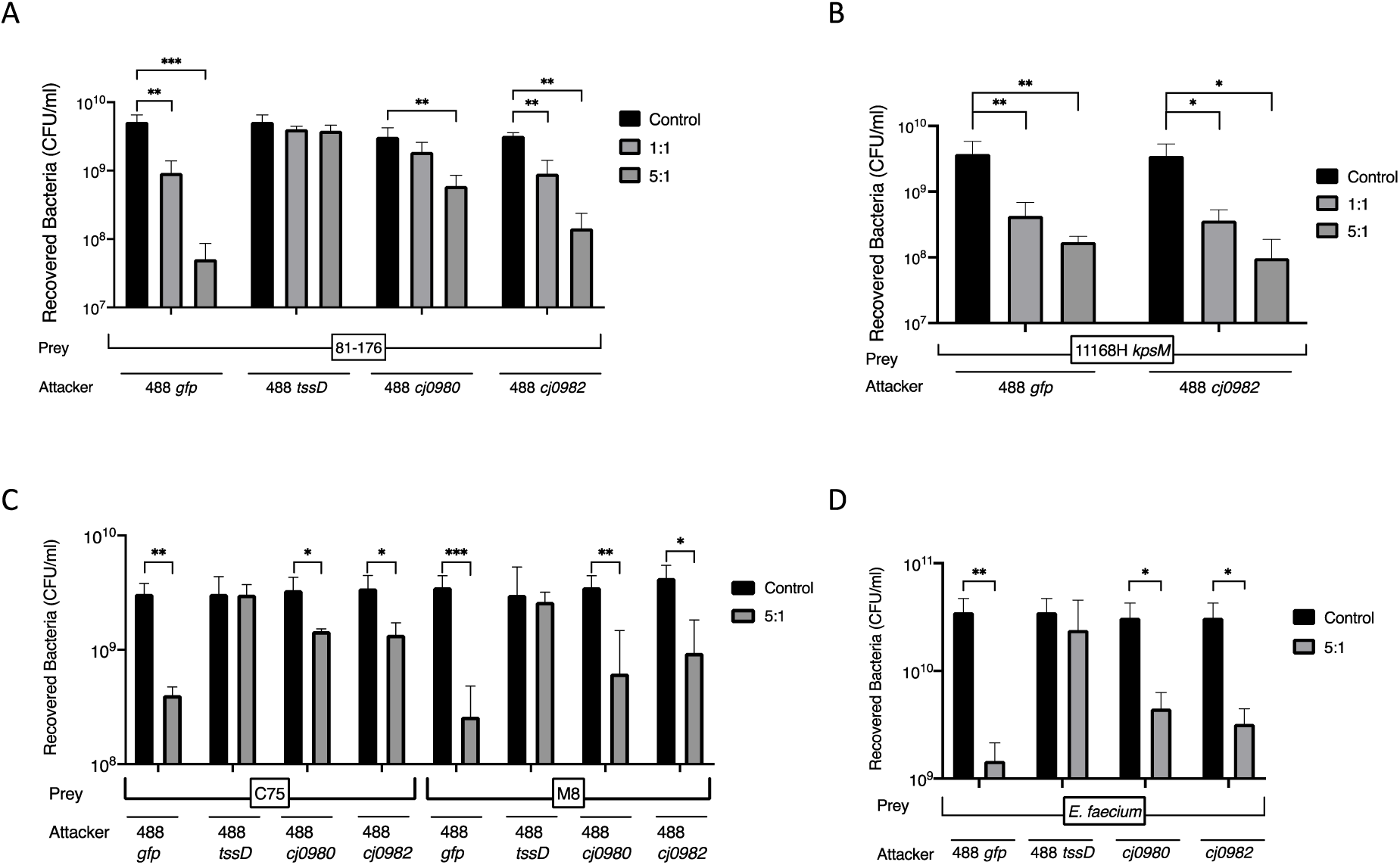
Recovered input and output prey CFU/ml following interspecies competition assays with the putative effector mutants. The indicated *C. jejuni* attackers (488 *gfp,* 488 *tssD* mutant, 488 *cj0980* mutant or 488 *cj0982* mutant) were mixed and incubated with prey in a 1:1 and/or 5:1 ratio of attacker to prey and co-incubated on *Brucella* agar for 16 hours under microaerobic conditions. After incubation bacteria were harvested, serially diluted, and plated onto selective agar to select for either the attacker or prey bacteria. The recovered input and output CFU/ml of the prey (A) 81-176 strain, (B) 11168H *kpsM* mutant, (C) *C. coli* C75 and *C. coli* M8 strains or (D) *E. faecium* strain are displayed and compared against the respective prey input control incubated in the absence of the attacker. Experiments were repeated in three biological replicates with three technical replicates. Asterisks denote a statistically significant difference (*= *p* < 0.05; ** = *p* < 0.01; *** = *p* < 0.001).

**Figure S9:**
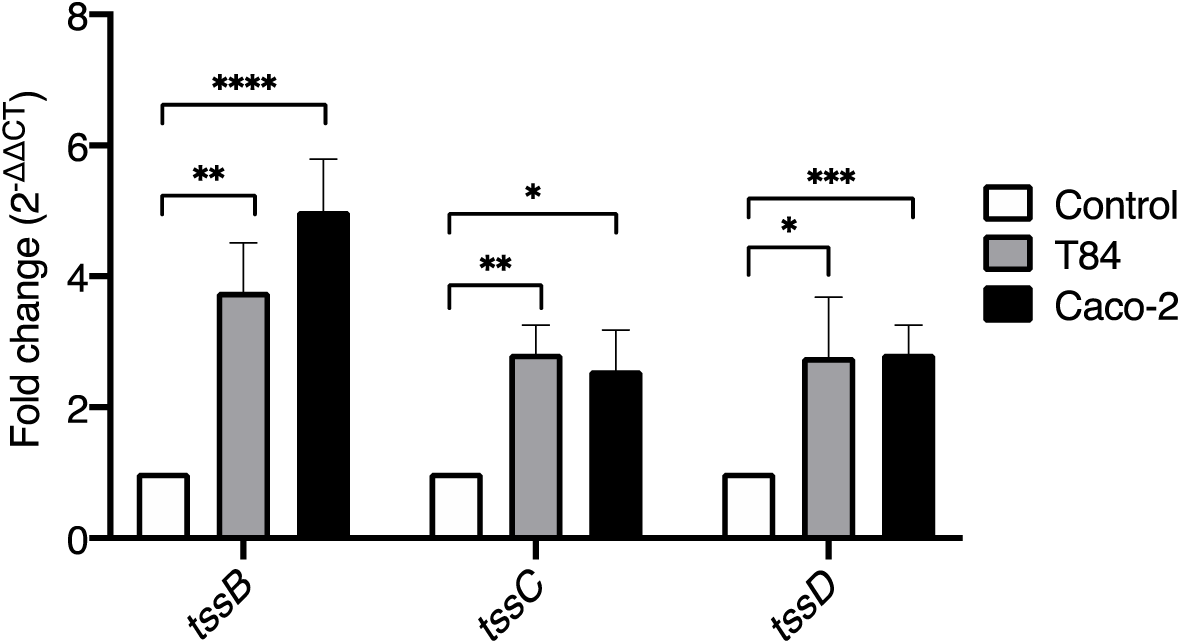
qRT-PCR analysis of T6SS gene expression in T84 and Caco-2 cells following infection by *C. jejuni* 488 wild-type strain. T84 and Caco-2 cells were infected with *C. jejuni* 488 wild-type strain for 3 hours at 37°C in a 5% CO_2_ incubator (MOI of 200:1). After infection, RNA was extracted, converted into cDNA which was used as a template for qRT-PCR analysis. The expression of *tssB*, *tssC* and *tssD* was analysed with fold change comparisons made against the expression levels of a cell monolayer-free *C. jejuni* 488 wild-type strain control incubated under the same conditions. *rpoA* was used an internal and experiments were repeated in three biological replicates with three technical replicates. Asterisks denote a statistically significant difference (*= *p* < 0.05; ** = *p* < 0.01; *** = *p* < 0.001; *** = *p* < 0.0001).

**Table S1.** *C. jejuni* strains used in this study.

| *C. jejuni* strain | Description | Antibiotic selection | Source |
| --- | --- | --- | --- |
|  | T6SS-positive strains |  |  |
| 488 | Brazilian human clinical isolate | N/A | (Liaw et al., 2019, Davies et al., 2019) |
| 488 *tssD* | Non-functional T6SS mutant constructed by insertion mutagenesis with a Kan<sup>r</sup> cassette disrupting *tssD*. | Kanamycin | Constructed by Dr. Janie Liaw (Liaw et al., 2019)<br>Obtained from the London School of Hygiene and Tropical Medicine<br>Campylobacter Resource Facility |
| 488 *tssD*<sup>+</sup> | Functional T6SS *tssD* mutant complemented with a copy of *tssD* using a pCJC1 plasmid. | Kanamycin and Chloramphenicol | Constructed by Dr. Janie Liaw (Liaw et al., 2019).<br>Obtained from the London School of Hygiene and Tropical Medicine<br>Campylobacter Resource Facility |
| 488 *cj0980* | Functional T6SS *cj0980* putative effector mutant constructed by insertion mutagenesis with a Kan<sup>r</sup> cassette disrupting *cj0980*. | Kanamycin | This study |
| 488 *cj0982* | Functional T6SS *cj0982* putative effector mutant of constructed by insertion mutagenesis with a Ery<sup>r</sup> cassette disrupting *cj0982*. | Erythromycin | This study |
| 488 *gfp* | Functional T6SS 488 wild-type strain expressing *gfp* reporter gene constructed by insertion mutagenesis with a pCJC1 plasmid harbouring *gfp*. | Chloramphenicol | This study |
| T6SS-negative strains |  |  |  |
| 11168H *kpsM* | Capsular polysaccharide mutant of the hypermotile variant of NCTC11168 wild-type strain. Constructed by insertion mutagenesis with a Kan<sup>r</sup> cassette disrupting *KpsM*. | Kanamycin | Constructed by Dr. Andrey V. Karlyshev (Karlyshev et al., 2000). Obtained from the London School of Hygiene and Tropical Medicine Campylobacter Resource Facility |
| 81-176 | Human clinical isolate from a case of gastroenteritis associated with raw milk consumption in the USA. | Tetracycline | (Korlath et al., 1985) |

**Table S2.** Bacterial prey strains and *E. coli* strains used in this study.

| Strain | Description | Source |
| --- | --- | --- |
| *C. coli* C75 | Tetracycline resistant<br>Vietnamese strain isolated from<br>farmed chicken faeces. | (Lehri et al., 2023) |
| *C. coli* M8 | Tetracycline Vietnamese strain<br>isolated from retail chicken<br>meat. | (Lehri et al., 2023) |
| *E. faecium* NCTC 12202 | Vancomycin-resistant<br>*Enterococcus* (VRE)<br>strain isolated from hospitalised<br>patient. | NCTC at the UK Health<br>Security Agency culture<br>collections (Uttley et al., 1989) |
| *E. coli* DH5α | Commercially available strain<br>used for competitive index<br>assays. | New England BioLabs (NEB;<br>UK) |
| XL1-Blue<br>Competent<br>Cells | Competent *E. coli* used for<br>transformation | Agilent Technologies |

**Table S3.**
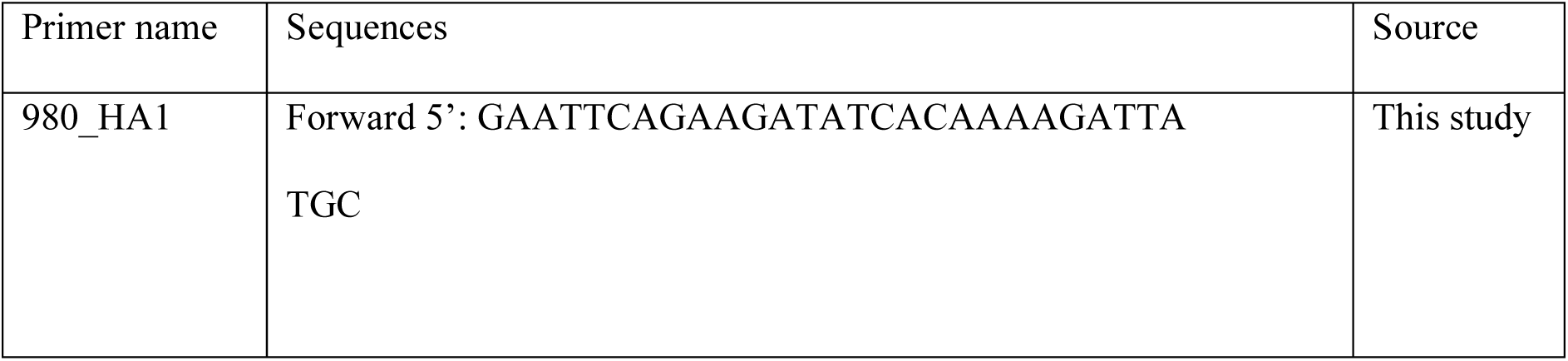
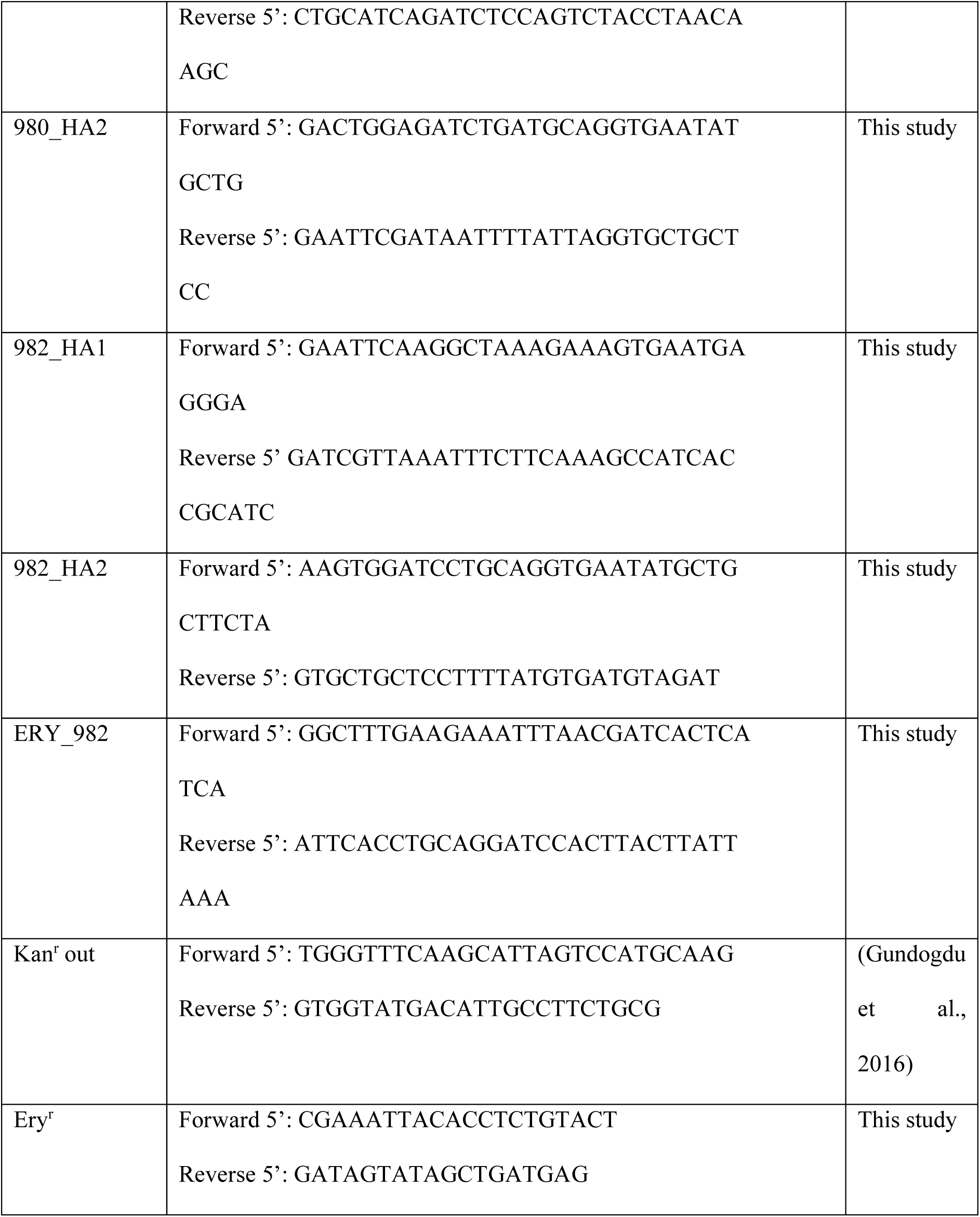
Mutagenesis primers used in this study.

**Table S4.**
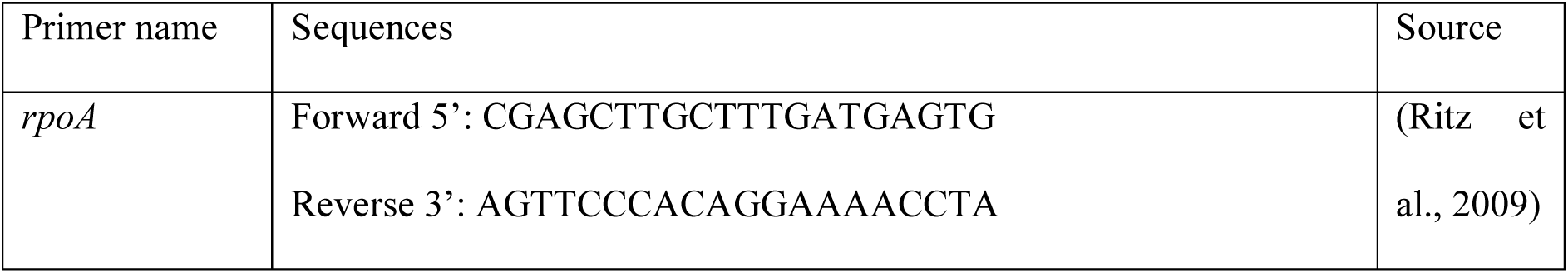
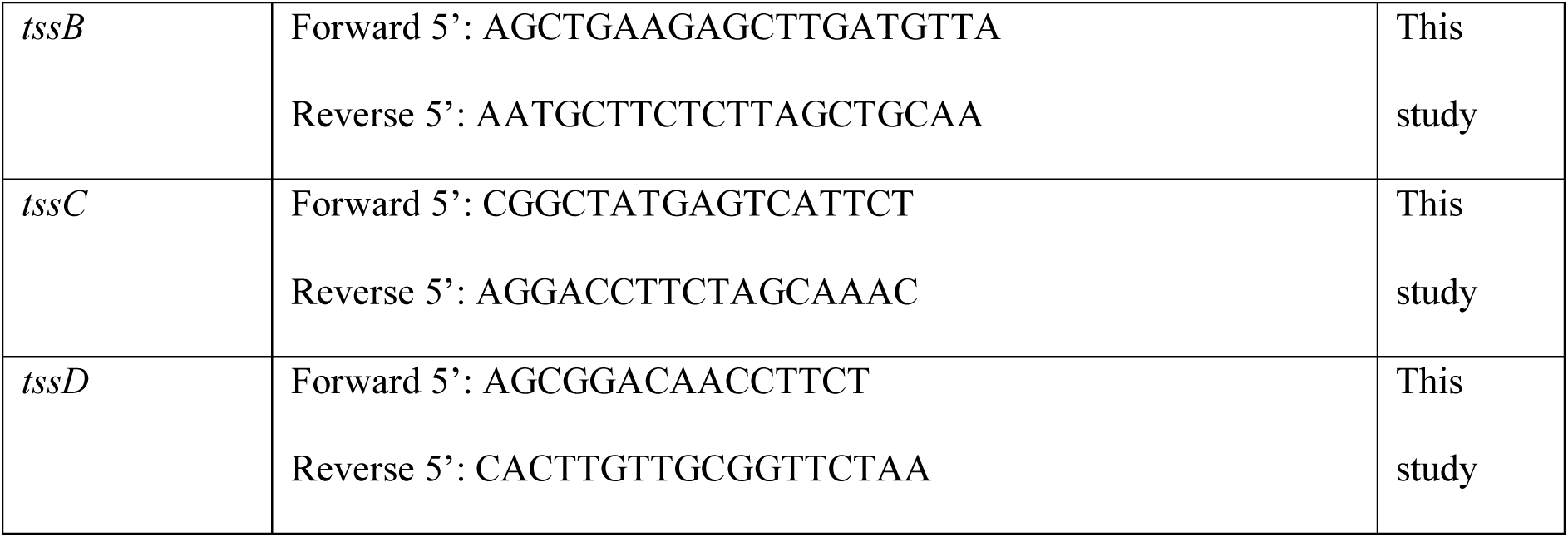
Oligonucleotide primers used in this study.

**Table S5.** Plasmids used during this study.

| Plasmid name | Description | Source |
| --- | --- | --- |
| pGEM®-T Easy | Commercially available cloning vector | Promega (UK) |
| pCJC1 | Complementation vector harbouring *gfp* and Cam<sup>r</sup> | (Jervis et al., 2015) |

